# Targeting tankyrase scaffolding in the β-catenin destruction complex by PROTAC overcomes the limitation of catalytic inhibitors in cancer

**DOI:** 10.1101/2025.09.22.677768

**Authors:** Qian Wang, Liping Li, Lin You, Shuai Wang, Lei Han, Bingnan Wang, Liping Yao, Yesu Addepalli, Yong Lu, Ilgen Mender, Ann M. Flusche, Chiho Kim, Nageswari Yarravarapu, Andrew Lemoff, Lawrence Lum, Jerry W. Shay, Yonghao Yu, Chuo Chen

## Abstract

Aberrant WNT/β-catenin signaling drives tumorigenesis and metastasis in cancer. Although enzymatic inhibitors of tankyrase (TNKS) effectively block AXIN degradation and stabilize the β-catenin destruction complex (DC), they have demonstrated limited efficacy in various cancer models. Here we demonstrate that, unexpectedly, the induction of AXIN puncta represents a major barrier to achieving therapeutic efficacy. Mechanistically, catalytic inhibition of TNKS prevents TNKS turnover and drives its accumulation in the DC, wherein the scaffolding function of TNKS induces AXIN puncta formation, rigidifies the DC, and impedes β-catenin turnover. Chemically induced degradation of TNKS overcomes this limitation by stabilizing AXIN without puncta formation, providing a deeper suppression of the WNT/β-catenin pathway activity and the proliferation of colorectal cancer cells harboring dysfunctional APC mutations. Collectively, these findings provide an explanation for the unsatisfactory outcomes of drugging the WNT/β-catenin signaling pathway by TNKS inhibitors and highlight TNKS degradation as a promising approach to treat WNT/β-catenin-driven cancers.

## Introduction

WNT/β-catenin signaling plays an important role in cell growth, differentiation, and migration^1,2^. Dysregulation of this pathway enables reprogrammed cancer cells to proliferate, metastasize and resist chemo- and radiotherapies. In colorectal cancer (CRC), loss-of-function mutations in adenomatous polyposis coli (APC) are the major drivers of aberrant WNT/β-catenin signaling^3^. However, therapeutic targeting this oncogenic pathway remains unsuccessful despite extensive efforts^4^. For example, although silencing tankyrase (TNKS1/2, also known as PARP5A/B and ATRD5/6 and encoded by *TNKS* and *TNKS2*, collectively referred to as TNKS hereafter)^5–7^ phenocopied APC restoration and prevented tumorigenesis in *APC*-null mice^8^, TNKS inhibitors (TNKSi) failed to show promising efficacy in various in vitro and in vivo CRC models. Addressing the discrepancy between the genetic and pharmacological interception of WNT/β-catenin signaling through TNKS may provide a path toward developing new CRC treatments.

The axis inhibition protein (AXIN) and APC are both scaffolding proteins of the β-catenin destruction complex (DC) that likely exists as biomolecular condensates in the cytoplasm to prime β-catenin for proteasomal degradation^9–11^. As TNKS-catalyzed poly(ADP-ribose)-dependent ubiquitination (PARdU) is required for the turnover of AXIN^12–14^, blocking the enzymatic function of TNKS by IWR1 or XAV939 leads to AXIN accumulation^12,15^, thereby enhancing the ability of the DC to promote β-catenin degradation. The accompanied AXIN puncta formation is generally attributed to AXIN accumulation and is considered to play a positive role in suppressing WNT/β-catenin signaling. However, TNKS also associates with the DC^16–18^, where its contributions to the DC function remains poorly understood. Specifically, because TNKS regulates its own abundance through self-poly(ADP-ribosylation) (PARylation), catalytic inhibition also promotes drastic TNKS accumulation^12^. It is unclear whether this feedback mechanism affects the DC function and results in the limited anticancer efficacy of TNKSi, as inhibitor-induced TNKS accumulation may support WNT/β-catenin signaling through molecular scaffolding independently of its catalytic function. Indeed, deletion of the catalytic PARP domain of TNKS attenuated, but did not fully suppress the WNT-induced transcription of the TCF/LEF-controlled genes^12,16,19^. By contrast, removal of the sterile alpha motif (SAM) domain or the ankyrin repeat clusters (ARCs) of TNKS abolished the WNT/β-catenin pathway activity more completely.

To understand the role of TNKS scaffolding in WNT/β-catenin signaling, we developed a TNKS-targeting proteolysis-targeting chimera (PROTAC)^20,21^ as a selective TNKS degrader (TNKSd). Comparison between TNKSd and TNKSi allowed us to understand the catalysis-independent scaffolding function of TNKS and uncover the mechanism underlying the ineffectiveness of TNKSi against cancers. We found that inhibiting TNKS by TNKSi led to the accumulation of TNKS within the DC, which induced AXIN puncta formation, promoted maturation^22,23^ of the DC condensates, and impeded β-catenin turnover. In contrast, degrading TNKS by TNKSd enriched AXIN without inducing puncta or impairing the dynamics of the DC. As a result, TNKSd provided deeper suppression of the WNT/β-catenin pathway activity as well as better control of the proliferation of CRC cells harboring dysfunctional APC mutations. Collectively, our study provides new insights into the roles of TNKS in the DC and presents a promising strategy to enhance the therapeutic efficacy of TNKS-targeting anticancer treatments.

## Results

### IWR1-POMA induces TNKS degradation

To study the role of TNKS scaffolding, we first developed a PROTAC molecule to enable selective degradation of TNKS1 and TNKS2 at the same time by small molecules. Compared to conventional genetic approaches, this chemical approach helps address gene redundancy more conveniently and effectively. TNKS uses nicotinamide adenine dinucleotide (NAD^+^) as the ADP-ribose donor. The corresponding pocket in the PARP domain contains two discrete sites, one for nicotinamide (NI) and the other for adenosine (AD) binding. IWR1 is a highly selective TNKSi that exploits the AD site unique to TNKS, and IWR6 is a promiscuous but potent TNKSi structurally similar to XAV939 that binds to the NI site common to all PARPs^12,24^. Based on the crystal structures of TNKS1 and TNKS2 in complex with IWR1 or XAV939 (Figure S1), we envisioned that modifying the quinoline group of IWR1 and the phenyl group of IWR6 would not negatively impact TNKS-binding. We have thus created a small library of 30 PROTAC molecules by attaching pomalidomide^25^ or VH032^26^ through a PEG chain of various lengths to IWR1 or IWR6 (Figure S2A) for engaging cereblon (CRBN) or the von Hippel‒Lindau tumor suppressor protein (VHL), respectively, and examined their ability to induce TNKS degradation.

To facilitate the discovery of a TNKSd, we also developed a luciferase assay to measure the level of TNKS1 in a high-throughput manner. Using the CRISPR-assisted insertion tagging (CRISPaint) technology^27^, we introduced a nanoluciferase (NanoLuc) tag to the C-terminus of TNKS1 in HAP1 cells (Figure S2B). We chose to monitor TNKS1 because it is constitutively expressed while TNKS2 is expressed minimally under basal conditions. By following the luciferase activity, the knockdown efficiency of all 5 series of the PROTAC molecules could be quantified easily and accurately using a robotic liquid dispenser system in 96-well plates, which significantly improved the efficiency over the traditional method that relies on Western blotting and image processing. Using this luciferase assay, we found that the IWR1-based PROTACs eliminated TNKS1 significantly more effectively than the IWR6-based PROTACs (Figure S2C). Additionally, CRBN mediated TNKS1 degradation more efficiently than VHL. Inserting a triazole group between the quinoline group and the PEG chain further enhanced the degradation efficiency. However, replacing the PEG linker with a nonpolar or rigid linker did not improve the efficacy, and employing G007-LK^28^ as the warhead with various linker strategies led to reduced potency and degree of degradation (Figure S2D).

IWR1-TP4-Poma (IWR1-POMA hereafter) is the most potent PROTAC among these series. It initiated TNKS1 degradation at 1 nM, operated with a DC_50_ value of 60 nM, and reached 96% degradation at 1.2 μM in HAP1 cells (Figure 1A). Western blot analysis confirmed that IWR1-POMA could also deplete TNKS1 and suppress the induction of TNKS2 in a dose and time-dependent manner in DLD-1 CRC cells that carry truncating APC mutations (Figure 1B and C). The level of TNKS did not recover 36 hours after IWR1-POMA was removed, indicating a durable drug effect (Figure S3A). Addition of IWR1, pomalidomide, or MG132 blocked TNKS degradation (Figure S3B and C), supporting that IWR1-POMA functioned through the designed mode of action. The generality of IWR1-POMA to induce TNKS degradation was demonstrated in SW480, HT-29 and HeLa cells (Figure S3D). The ability of IWR1-POMA to stabilize AXIN and promote β-catenin degradation in the cytoplasm was confirmed further in HEK293 cells (Figure 1D). Compared to IWR1, IWR1-POMA provided better control of the cytosolic β-catenin level. The performance of this PROTAC molecule is particularly notable, considering that shutting down the catalysis of TNKS leads to a >30-fold accumulation of total TNKS, creating a nearly 3-orders-of-magnitude difference in the levels of TNKS between catalytic inhibition and chemically induced degradation. Although hook effect was observed at high concentrations, presumably due to the formation of nonproductive binary protein-ligand complexes under these conditions, to our knowledge, this is the first example of targeted degradation of an auto-regulated protein.

**Figure 1.**
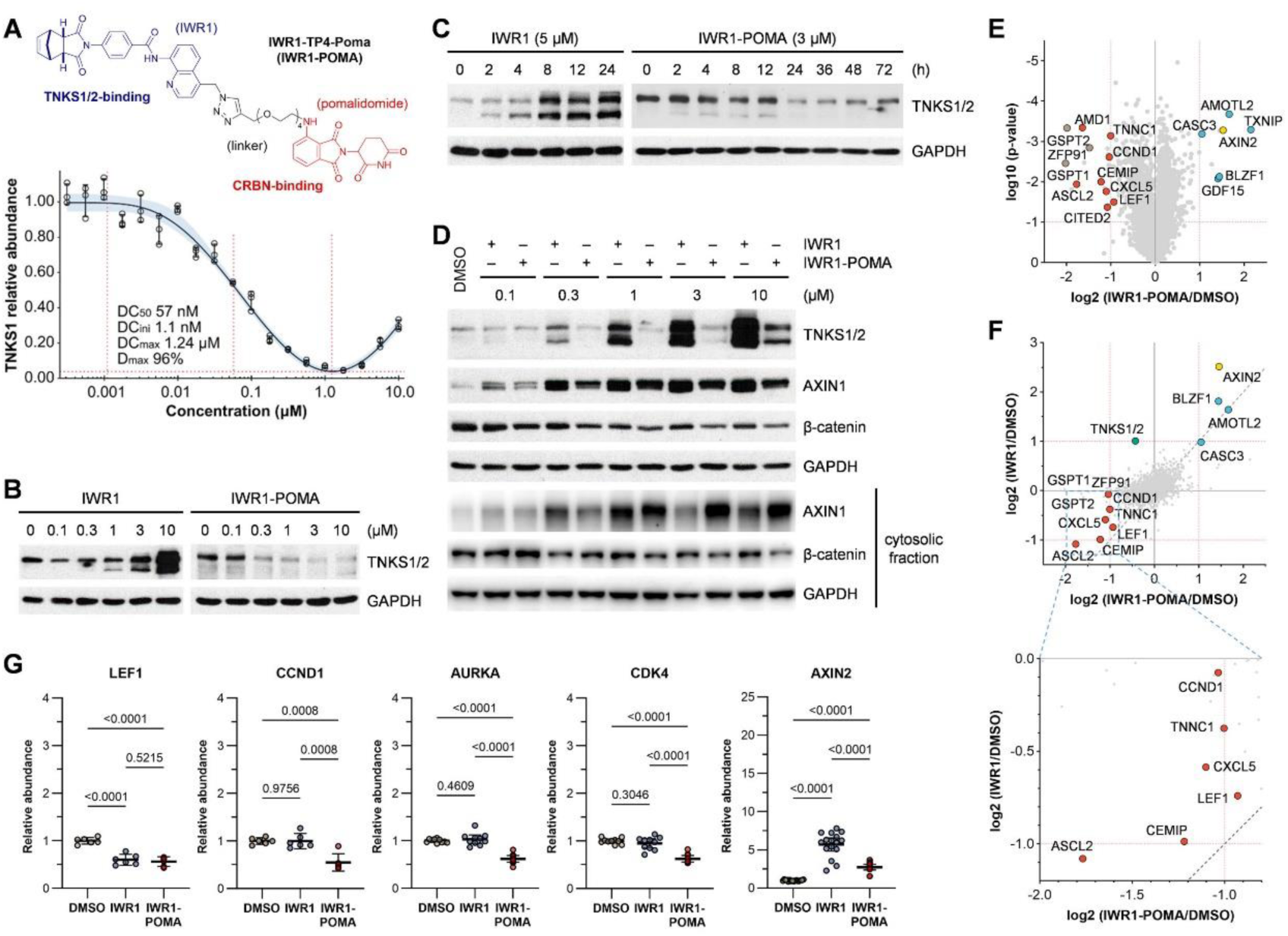
Characterization of IWR1-POMA. (A) IWR1-POMA induced TNKS degradation with a DC50 value of 60 nM and reached a nearly complete depletion of TNKS1 at 1.2 µM in HAP1 cells. The dose-response curve is presented as mean ± SEM (n = 3 biological samples) with 95% confidence interval (CI) indicated by the shaded area. (B and C) IWR1 induced massive TNKS accumulation while IWR1-POMA promoted deep degradation in a dose and time-dependent manner in DLD-1 cells. (D) IWR1-POMA promoted a more complete degradation of β-catenin than IWR1 in HEK293 cells cultured with Wnt3A conditioned media. (E) Volcano plot of the TMT experiment results. TNKS substrates (light blue) accumulated and WNT/β-catenin-regulated proteins (red) downregulated in DLD-1 cells treated with IWR1-POMA (3 μM). AXIN2 (yellow) is both a TNKS substrate and a WNT target. GSPT1/2 and ZFP91 (light brown) are the only off-targets identified. (F) Comparative analysis of the TMT experiment showing that the levels of several WNT/β-catenin regulated proteins (red) were preferentially reduced by IWR1-POMA relatively to IWR1 treatment. The dashed line denotes equal protein abundance between the two treatments. Data points above this line in Quadrant III represent proteins selectively downregulated by IWR1-POMA relatively to IWR1. (G) Relative peptide abundances of selected WNT targets in the TMT experiment with p-values calculated by two-tailed unpaired t-test.

To confirm the ability of IWR1-POMA to engage TNKS, we prepared IWR1-PEG3-NCT as a bioluminescence resonance energy transfer (BRET) tracer (Figure S3E)^29^. We then treated HAP1-TNKS1-NanoLuc cells with MG132 to increase the signal-to-noise ratio and to normalize the TNKS level in the presence of IWR1 and IWR1-POMA. Addition of IWR1 effectively reduced the BRET signals; however, the intrinsic fluorescence^30^ of IWR1-POMA prevented the assessment of its binding affinity by this method. Nonetheless, the observation of significant BRET signals of IWR1-POMA at high concentrations prompted us to test whether it could be used directly as a BRET acceptor for NanoLuc despite poor spectral overlap^31^. Indeed, the BRET signals increased in a dose-dependent manner after adding IWR1-POMA to HAP1-TNKS1-NanoLuc cells in the presence of MG132 (Figure S3F). Accordingly, we titrated IWR1 into HAP1-TNKS1-NanoLuc cell in the presence of 0.5 μM of IWR1-POMA and found that IWR1 suppressed the BRET signals with an IC_50_ value of 0.25 μM (Figure S3G). Repeating this experiment in 293T cells transfected with TNKS1-NanoLuc plasmid improved the BRET signal-to-noise ratio and confirmed that the pomalidomide chromophore can be used as a BRET pair for NanoLuc. It also confirmed that IWR1-POMA (0.5 μM) binds to TNKS1 with a slight reduction in affinity as compared to IWR1 (IC_50_ 0.15 μM).

To examine the degradation specificity of IWR1-POMA, we performed proteome-wide expression profiling on DLD-1 cells treated with DMSO, IWR1 or IWR1-POMA for 16 h. Using a tandem mass tag (TMT)-based, multiplexed quantitative mass spectrometric (MS) approach with two biologically independent replicate samples (Figure S4A), we identified 7,884 proteins, among which 5,822 could be quantified with high confidence (<1% FDR). Correlation analysis confirmed the consistency of the quantitative analysis of the two biological replicates (Figure S4B). However, accurate quantification of TNKS1 proved challenging, as only a low-abundance peptide corresponding to TNKS1 was detected in this experiment. Nonetheless, IWR1-POMA induced the accumulation of AXIN2 along with other TNKS substrates (Figure 1E). LEF1 and several other WNT targets were also effectively downregulated, confirming the on-target drug effects. Pairwise binary comparison of the drug treatment versus control indicated better control of WNT signaling by IWR1-POMA in addition to high degradation specificity (Figure 1F and G). GSPT1/2 and ZFP91, three common off-targets of the IMiD-based PROTACs, were the only perturbations not related to the WNT pathway. None of the other 7 PARPs and 81 NAD^+^/NADP^+^-dependent enzymes identified in this MS experiment were significantly affected by IWR1-POMA (Figure S4C).

### PROTAC suppresses catalysis-independent WNT/β-catenin signaling by TNKS

A previous study showed that polymerization of PARylation-incompetent TNKS can drive WNT/β-catenin signaling^19^. To understand whether the lack of efficacy of TNKSi in suppressing CRC cell growth originated from the PARylation-independent WNT/β-catenin signaling induced by the accumulated TNKS upon catalytic inhibition, we treated Wnt3A-stimulated 293T cells with IWR1 or IWR1-POMA. We reason that, by removing both the catalysis-dependent and independent functions of TNKS, IWR1-POMA may suppress WNT/β-catenin signaling more completely than IWR1 that inhibits AXIN PARylation but also promotes TNKS accumulation and polymerization. Indeed, IWR1-POMA promoted β-catenin degradation significantly more effectively than IWR1 under various doses of Wnt3A (Figure S5A). To investigate the individual contribution of TNKS1 and TNKS2 to WNT/β-catenin signaling, we co-transfected a low dose of TNKS1 plasmid together with SuperTopFlash (STF), a luciferase reporter for the WNT/β-catenin activity, into 293T-TNKS1/2-DKO cells wherein both TNKS1 and TNKS2 were deleted by CRISPR^32^ (Figure S5B). We found that addition of IWR1 suppressed the TNKS1-induced STF activity but not completely (Figure 2A and S5C). There was a small but noticeable residual activity, which could not be removed by increasing the concentration of IWR1. In contrast, IWR1-POMA induced TNKS1 degradation and promoted deeper suppression of the STF activity. Similarly, IWR1-POMA reduced the level of TNKS2 and provided better control of the STF activity than IWR1 in TNKS1/2-DKO cells expressing TNKS2 (Figure 2B and S5D). As such, both TNKS1 and TNKS2 can contribute to WNT signaling that persists even in the presence of catalytic inhibition of TNKS PARylation.

**Figure 2.**
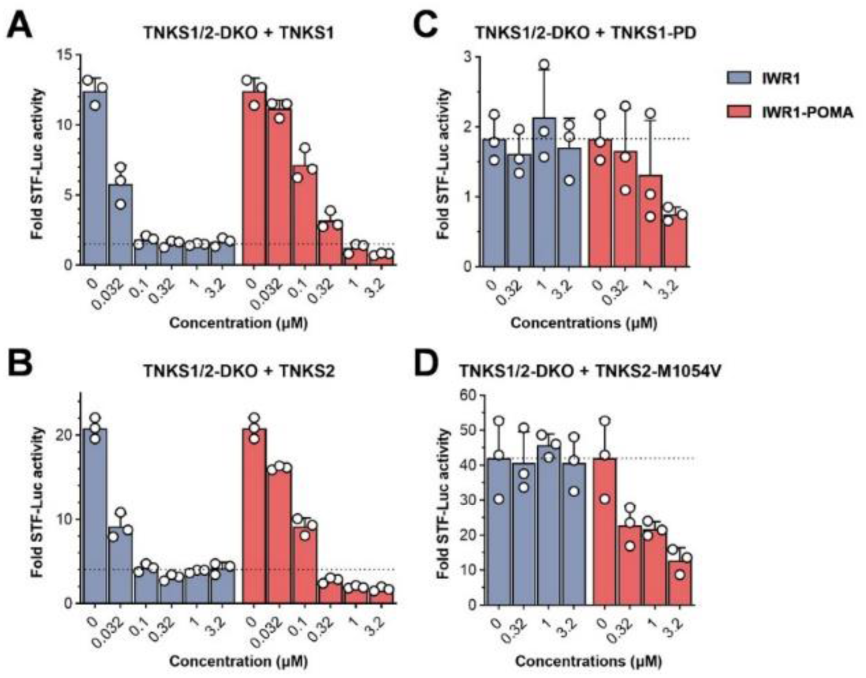
IWR1-POMA degraded TNKS to suppress both catalysis-dependent and independent WNT signaling. (A) 293T-TNKS1/2-DKO cells were transfected with FLAG-TNKS1 and STF plasmids and then treated with IWR1 or IWR1-POMA. In contrast to IWR1 that plateaued in suppressing the WNT/β-catenin signaling induced by TNKS1, IWR1-POMA allowed for a more complete control of the pathway activity. (B) 293T-TNKS1/2-DKO cells transfected with FLAG-TNKS2 and STF plasmids and then treated with IWR1 or IWR1-POMA similarly showed that catalytic inhibition led to residual WNT/β-catenin activity. (C) 293T-TNKS1/2-DKO cells were transfected with 3×FLAG-TNKS1-PD and STF plasmids and then treated with IWR1 or IWR1-POMA. IWR1-POMA suppressed the luciferase activities whereas IWR1 had no effect. (D) 293T-TNKS1/2-DKO cells transfected with FLAG-TNKS2-M1054V and STF plasmids and then treated with IWR1 or IWR1-POMA. TNKS2 exhibited robust scaffolding effects. All data are presented as mean ± SEM (n = 3 biological samples).

We next used PARylation-incompetent variants of TNKS to verify whether TNKS1 and TNKS2 can promote WNT/β-catenin signaling independently of their catalytic function. The catalytic site of TNKS is highly conserved. Deactivation of the H-Y-E triad of TNKS1 with H1184A and E1291A mutations^33^ abolishes the binding of NAD^+^ and the accepting ADP ribose. This PARP-dead (PD) variant induced notable WNT/β-catenin signaling in the TNKS1/2-DKO cells even at low doses, confirming that TNKS1 has a catalysis-independent functional role in the DC (Figure 2C and S5E). Addition of IWR1-POMA degraded TNKS1-PD and reduced the pathway activity, whereas treatment with IWR1 had no effect. Similarly, TNKS2-M1054V having an impaired ability to interact with the accepting ADP ribose^34^ induced robust WNT/β-catenin signaling and was insensitive to IWR1 treatment (Figure 2D and S5F). In contrast, IWR1-POMA degraded this catalytically inactive variant of TNKS2 and suppressed the corresponding STF activity in a dose-dependent manner. Collectively, these data support the hypothesis that both TNKS1 and TNKS2 play an additional role in the DC to regulate WNT/β-catenin signaling through a mechanism beyond catalytic PARylation of AXIN^19^. Degradation of TNKS eliminates both the enzymatic and non-enzymatic activities of TNKS, providing better control of the WNT/β-catenin signaling than catalytic inhibition (Figure S5G).

### Tankyrase controls the dynamic assembly of the DC

AXIN interacts with APC to work as a molecular scaffold for the DC to catalyze β-catenin phosphorylation and ubiquitination. TNKS also associates with this complex^16–18^ to control the AXIN level. Although much is known about the catalytic function of TNKS, its scaffolding function^19,35^ is poorly understood. To study the non-enzymatic function of TNKS in WNT signaling, we transfected 293T cells with a low dose of AXIN1-mCherry plasmid and confirmed that TNKS colocalized with AXIN1 to form micrometer-sized puncta (Figure 3A and S6). In contrast, in TNKS1/2-DKO cells that lack TNKS, AXIN1-mCherry distributed diffusely throughout the cytoplasm as small aggregates (Figure 3B). However, AXIN puncta formed readily when AXIN1-mCherry was expressed together with TNKS1 in the TNKS1/2-DKO cells (Figure 3C). Similarly, TNKS2 also induced AXIN puncta in TNKS1/2-DKO cells (Figure 3D), suggesting that TNKS1 and TNKS2 play a redundant structural role in the DC to support AXIN puncta formation. To further verify whether AXIN PARylation is involved in the puncta formation, we expressed AXIN1-mCherry together with catalytically inactive variants of TNKS and found that both TNKS1-PD and TNKS2-M1054V could induce AXIN puncta effectively (Figure 3E and F). Quantitative imaging analysis confirmed that TNKS1 and TNKS2 promoted AXIN puncta formation through molecular scaffolding independently of their ability to catalyze protein PARylation (Figure 3G).

**Figure 3.**
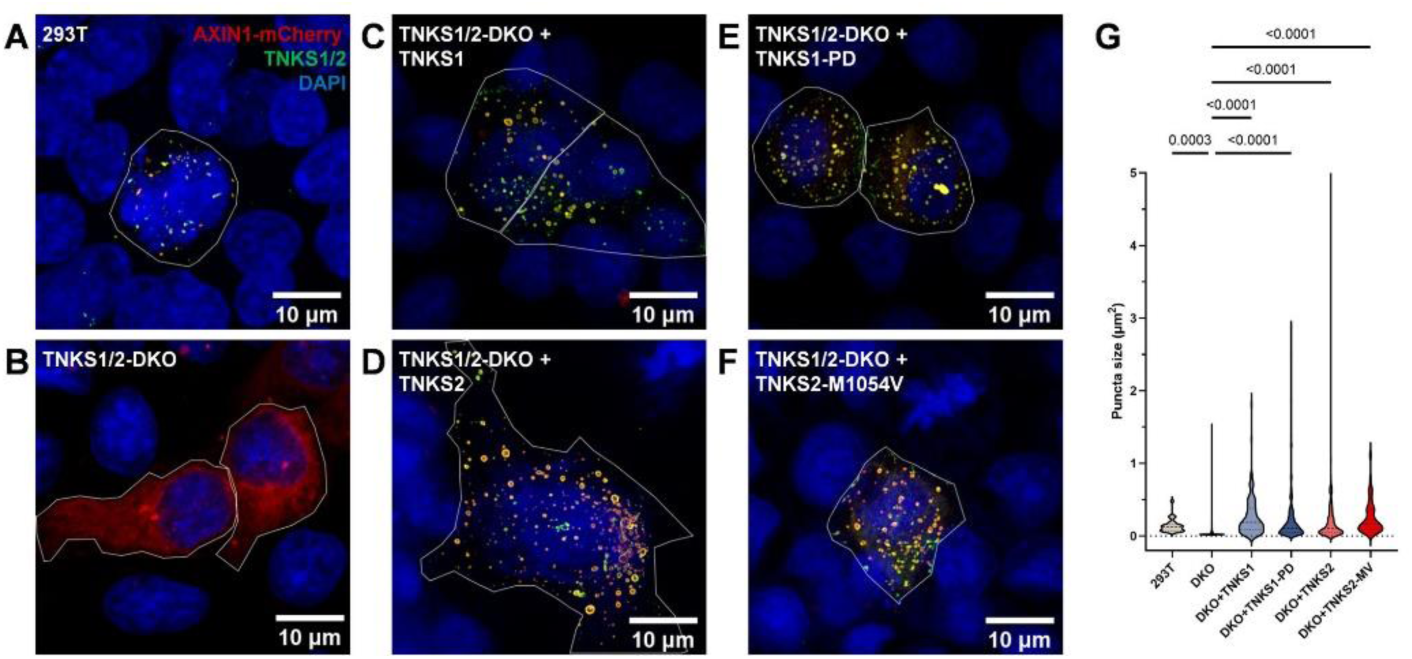
TNKS is required for AXIN puncta formation. (A) AXIN colocalized with TNKS and formed puncta in 293T cells transfected with AXIN1-mCherry plasmid. (B) AXIN distributed diffusely throughout the cytoplasm in TNKS1/2-DKO cells transfected with AXIN1-mCherry plasmid. (C) Introduction of TNKS1 restored AXIN puncta in TNKS1/2-DKO cells transfected with AXIN1-mCherry and TNKS1 plasmids. (D) TNKS2 also induced puncta formation in TNKS1/2-DKO cells transfected with AXIN1-mCherry and TNKS2 plasmids. (E) The catalytic function of TNKS1 is not required for puncta formation as demonstrated in TNKS1/2-DKO cells transfected with AXIN1-mCherry and TNKS1-PD plasmids. (F) Catalytically inactive TNKS2-M1054V also promoted AXIN puncta formation effectively in TNKS1/2-DKO cells transfected with AXIN1-mCherry and TNKS2-M1054V plasmids. (G) Quantification of the AXIN1-mCherry puncta in 293T cells (n = 28 biological samples), TNKS1/2-DKO cells (n = 230 biological samples), TNKS1/2-DKO cells in the presence of TNKS1 (n = 115 biological samples), TNKS1/2-DKO cells in the presence of TNKS2 (n = 229 biological samples), TNKS1/2-DKO cells in the presence of TNKS1-PD (n = 246 biological samples), and TNKS1/2-DKO cells in the presence of TNKS2-M1054V (n = 86 biological samples). Data are presented with p-values calculated by two-tailed unpaired t-test.

Both IWR1 and IWR1-POMA stabilize AXIN to suppress WNT/β-catenin signaling. However, IWR1 induces TNKS accumulation while IWR1-POMA promotes TNKS degradation. Given the observation that AXIN puncta formation requires TNKS, we next examined how these TNKS modulators affect the assembly of the DC. As expected, treating SW480 cells with IWR1 induced AXIN puncta while IWR1-POMA did not (Figure 4A and B). Similarly, IWR1 induced AXIN puncta in 293T cells wherein the endogenous AXIN1 was labeled with RFP at its C-terminus by CRISPR^36^, but IWR1-POMA failed to induce AXIN puncta (Figure 4C and D). As 293T cells harbor wild-type APC and SW480 cells carry a truncating APC without the AXIN-binding sites^37,38^, the ability of TNKSi to induce AXIN puncta is independent of AXIN-APC binding. These results also suggest that tagging AXIN1 with a fluorescent protein does not affect its ability to oligomerize.

**Figure 4.**
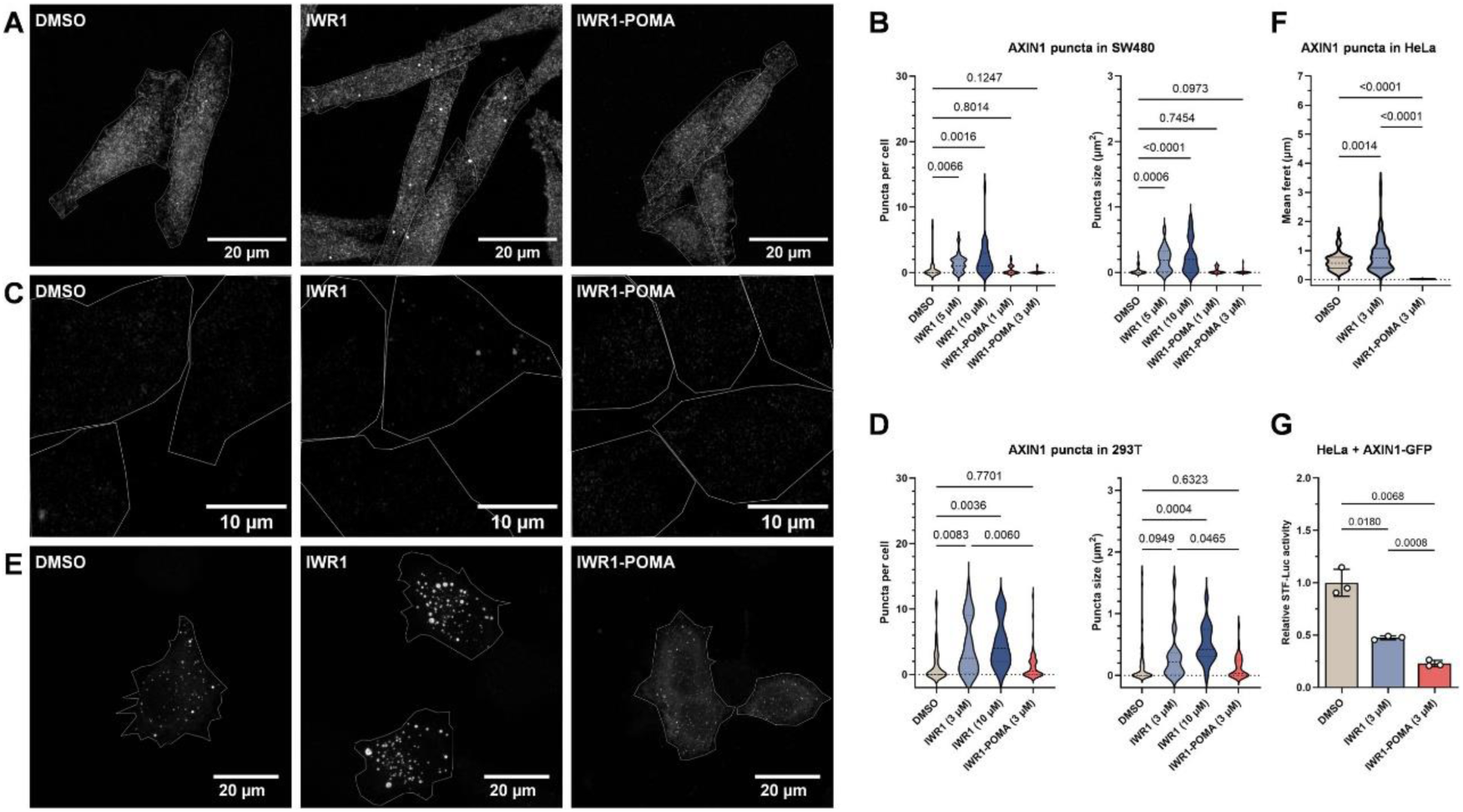
Chemically induced TNKS accumulation promoted AXIN puncta formation. (A) SW480 cells treated with DMSO, IWR1 (5 μM), or IWR1-POMA (1 μM) and stained with anti-AXIN1 antibody. (B) Numbers and sizes of the AXIN puncta in SW480 cells treated with DMSO (n = 36 biological samples), IWR1 (5 μM, n = 20 biological samples, or 10 μM, n = 31 biological samples), or IWR1-POMA (1 μM, n = 17 biological samples, or 3 μM, n= 24 biological samples) for 16 h. (C) 293T-AXIN1-dsRed-KI cells treated with DMSO, IWR1 (3 μM), or IWR1-POMA (3 μM). (D) Numbers and sizes of the AXIN puncta in 293T-AXIN1-dsRed-KI cells treated with DMSO (n = 82 biological samples), IWR1 (3 μM, n = 39 biological samples, or 10 μM, n = 32 biological samples), or IWR1-POMA (3 μM, n= 61 biological samples) for 16 h. (E) HeLa cells were transfected with AXIN1-GFP plasmid and then treated with DMSO, IWR1 (3 µM), or IWR1-POMA (3 µM). IWR1 promoted the formation of micrometer-sized AXIN puncta whereas IWR1-POMA dissolved them. (F) Numbers and sizes of the AXIN1 puncta in HeLa cells transfected with AXIN1-GFP plasmide and treated with DMSO (n = 88 biological samples), IWR1 (3 µM, n = 91 biological samples), or IWR1-POMA (3 µM, n = 151 biological samples) for 16 h. (G) HeLa cells transfected with AXIN1-GFP and STF plasmids and then treated with DMSO, IWR1 (3 μM) or IWR1-POMA (3 μM). IWR1-POMA suppressed WNT/β-catenin signaling significantly better than IWR1, indicating that AXIN puncta formation is not required for the DC to promote β-catenin degradation. Data are presented as mean ± SEM (n = 3 biological samples) with p-values calculated by two-tailed unpaired t-test for all data.

To further understand the functional significance of AXIN puncta, we transfected HeLa cells with a low dose of AXIN1-GFP plasmid. Again, IWR1 promoted the formation of micrometer-sized AXIN puncta that are significantly larger than those observed in the DMSO control sample (Figure 4E and F). In contrast, AXIN1 distributed diffusely throughout the cytoplasm when the cells were treated with IWR1-POMA. To determine the effects of AXIN puncta on WNT signaling, we expressed AXIN1-GFP together with STF in HeLa cells under the same conditions and found that IWR1-POMA suppressed the luciferase activity better than IWR1 (Figure 4G). Thus, AXIN puncta formation is not a prerequisite for the DC to catalyze β-catenin degradation as commonly believed. Instead, the large AXIN puncta induced by IWR1 are less functional DCs than the unaggregated AXIN complexes formed upon IWR1-POMA treatment. We have further confirmed that the N-terminally tagged AXIN1 and the C-terminally tagged AXIN2 also responded to IWR1 and IWR1-POMA treatments in the same manner (Figure S7C and D).

The DC has been proposed to exist as biomolecular condensates^9,10^. Consistently, AXIN1-GFP formed puncta that exhibited fusion behavior in U2OS cells (Figure S8), supporting the notion that the DC possess liquid-like properties, at least in the presence of excess AXIN1. TNKS interacts with both AXIN and APC and can self-aggregate to form insoluble filaments^17,34,35,39,40^. We suspected that the massive accumulation of TNKS induced by IWR1 rigidified the DC^22,23^ and limited its ability to turn over β-catenin. To test this hypothesis, we expressed AXIN1-mCherry and mNeonGreen-β-catenin in HeLa cells and then treated these cells with DMSO, IWR1 or IWR1-POMA. AXIN1-mCherry and mNeonGreen-β-catenin colocalized in all samples, indicating that labeling AXIN1 and β-catenin with a fluorescent protein did not disrupt the recruitment of β-catenin to the DC (Figure 5A). We then performed fluorescence recovery after photobleaching (FRAP) analysis on AXIN1-mCherry to investigate the structural role of TNKS in the DC. After photobleaching, the fluorescence signals of the AXIN-mCherry puncta in the DMSO control samples recovered quickly (*k* = 0.093 s^‒1^, n = 21), indicating a dynamic assembly of the DC (Figure 5B)^41,42^. However, the fluorescence signals of the AXIN1-mCherry puncta in the IWR1-treated sample recovered slowly (*k* = 0.036 s^‒1^, n = 16) and plateaued at 62% level, in contrast to 78% for the DMSO control, suggesting that IWR1 rigidified the DC. In contrast, the dynamics of the fluorescence recovery of the few very small AXIN1-mCherry puncta in the IWR1-POMA-treated sample was comparable to that of the DMSO control (*k* = 0.090 s^‒1^, 71% recovery, n = 21). Because both IWR1 and IWR1-POMA stabilized AXIN (Figure 5C), the rigidification of the DC by IWR1 is not a result of AXIN accumulation. Instead, as the DC puncta in the IWR1-treated sample was enriched with and that in the IWR1-POMA-treated sample depleted with TNKS, the reduction of the DC dynamics by IWR1 most likely originated from TNKS accumulation. We next performed FRAP analysis on mNeonGreen-β-catenin to examine the functional role of TNKS scaffolding in the DC using the dynamics of β-catenin as a measurement for its turnover rate (*k* = 0.024 s^−1^, 78% recovery, n = 11 for the DMSO control). In contrast to IWR1-POMA that accelerated the fluorescence recovery of mNeonGreen-β-catenin (*k* = 0.046 s^‒1^, 76% recovery, n = 10), IWR1 reduced the level of recovery by 20% without affecting the kinetics (*k* = 0.028 s^‒1^, 58% recovery, n = 16). These results suggest that TNKSi functions in part through cytosolic retention of β-catenin, and TNKS accumulation negatively impacts the exchange of β-catenin in the DC (Figure 5D). Collectively, our data support the hypothesis that TNKS scaffolding augments AXIN puncta to limit the ability of the DC to effectively degrade β-catenin.

**Figure 5.**
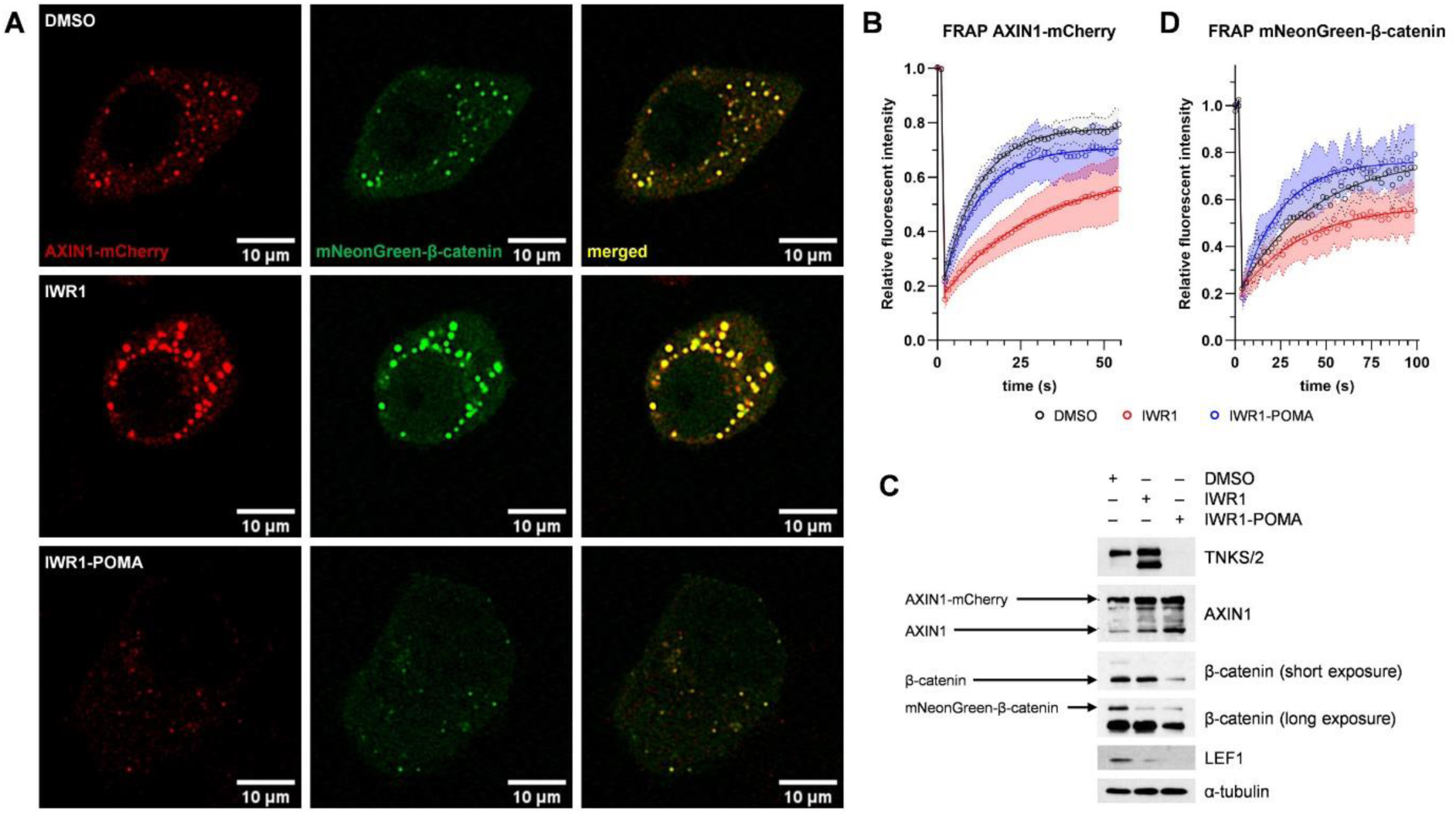
TNKS accumulation impeded the degradation of β-catenin. (A) HeLa cells were transfected with AXIN1-mCherry and mNeonGreen-β-catenin plasmids and then treated with DMSO, IWR1 (3 µM) or IWR1-POMA (3 µM). IWR1 promoted large AXIN1 puncta formation and IWR1-POMA dissolved the puncta. β-Catenin colocalized with AXIN1, indicating the proper assembly of the DC. (B) FRAP analysis provided support to the hypothesis that TNKS controls the dynamic assembly of the DCs. IWR1 treatment led to a slow recovery of the AXIN1-mCherry fluorescent signal after photobleaching. In contrast, IWR1-POMA treatment did not affect the mobility of AXIN1. The fluorescent recovery curves were fitted to one-phase association model. Data are presented as mean with 95% CI. (C) Western blot analysis of samples corresponding to Figure 5A confirmed the accumulation of TNKS by IWR1 and the depletion TNKS by IWR1-POMA. The deeper suppression of the Wnt/β-catenin signaling by IWR1-POMA is also supported by the reduced levels of total β-catenin. (D) FRAP analysis indicated that IWR1 limited the turnover of β-catenin. In contrast, IWR1-POMA accelerated the diffusion rate of β-catenin in the DC. The fluorescent recovery curves were fitted to one-phase association model. Data are presented as mean with 95% CI.

### Tankyrase degradation prevents CRC cell proliferation

Loss of functional APC impairs the DC function and results in constitutive WNT/β-catenin signaling that drives tumorigenesis and metastasis of CRC. Whereas depletion of both TNKS1 and TNKS2 by shRNA phenocopied APC restoration to prevent tumorigenesis in mice^8,43^, TNKSi only showed modest anti-proliferative activities in a limited number of CRC cell lines^44^. For example, IWR1 had minimal effects on the proliferation of DLD-1 cells that express truncated forms of APC. Remarkably, IWR1-POMA was able to suppress the formation of DLD-1 cell colonies under both normal and low serum conditions (Figure 6A and S9A). Similarly, the proliferation of SW480 and HT-29 cells carrying different truncating APC mutations were also better suppressed by IWR1-POMA (Figure 6A and S9B). The anti-proliferative activity also correlated with the ability of IWR1-POMA to better suppress WNT/β-catenin signaling in DLD-1 and SW480 cells (Figure S9C). Mechanistically, several WNT-controlled genes showed differential responses to IWR1 and IWR1-POMA. For example, whereas LEF1 was downregulated in both IWR1 and IWR1-POMA treated samples, cyclin D1 and Aurora kinase A responded to IWR1-POMA only in DLD-1 cells (Figure 1F and G and S9D). This observation is consistent with the ability of IWR1-POMA to better inhibit WNT/β-catenin signaling. Interestingly, IWR1-POMA also reduced the level of CDK4 while IWR1 had no effect, which may be explained by CDK4 being a direct target of c-MYC that in turn is a direct target of WNT/β-catenin. Indeed, IWR1-POMA reduced the level of c-MYC while IWR1 did not (Figure S9D). Taken together, targeted degradation of TNKS offers better control of the WNT/β-catenin pathway activities important to the maintenance of CRC cells than catalytic inhibition.

**Figure 6.**
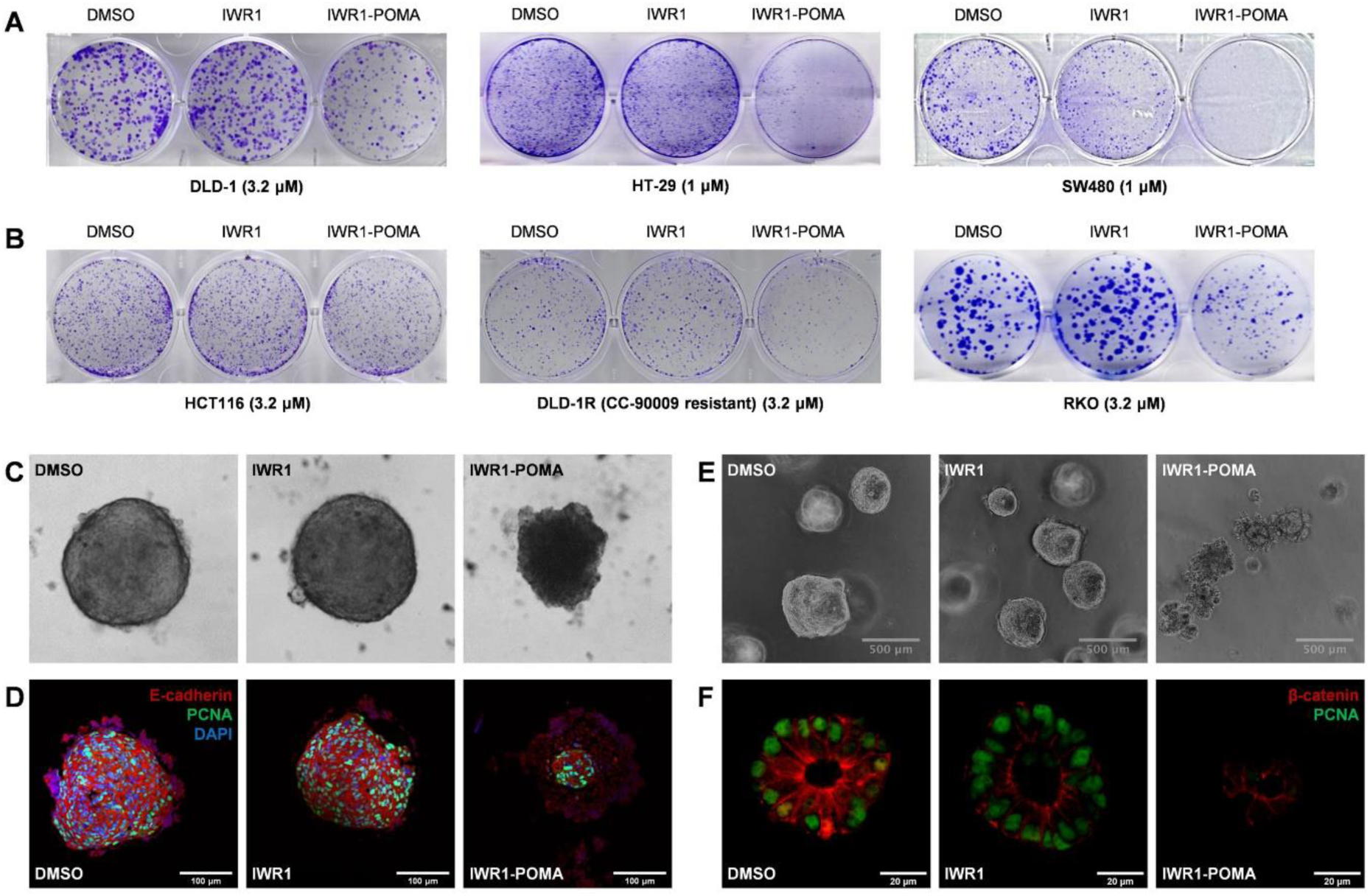
IWR1-POMA suppressed CRC cell growth. (A) IWR1-POMA suppressed the formation of DLD-1, SW480, and HT-29 CRC colonies significantly more effectively than IWR1. (B) HCT116 cells carrying a mutation in β-catenin that could not be processed by the DC were resistant to both IWR1 and IWR1-POMA. Additionally, DLD-1R cells obtained from cultivating DLD-1 cells with the GSPT1/2 inhibitor CC-90009 were still sensitive to IWR1-POMA treatment. IWR1-POMA could also inhibit RKO cells carrying full-length APC through the PTEN-AKT axis. (C) DLD-1 cells were developed into 3D spheroids and then treated with DMSO, IWR1 (5 µM), or IWR1-POMA (5 µM) for 10 d. The spheroids treated with IWR1-POMA lost the tight, spherical structure while IWR1 had no effect. (D) Immunostaining showed that the DLD-1 spheroids treated with IWR1-POMA only contained a small core of living cells while those treated with IWR1 remained highly proliferative. (E) IWR1-POMA (1 μM) suppressed the growth of PDM-7 patient-derived CRC organoids of approximately 200 μm in diameter with an apoptotic phenotype while IWR1 (1 μM) had no effect. (F) IWR1-POMA (1 μM) reduced the level of β-catenin and suppressed the proliferation of PDM-7 organoids grown from single cells significantly more effectively than IWR1 (1 μM).

GSPT1, a translation termination factor that binds eRF1 to mediate translation termination, is frequently identified as a neosubstrate of CRBN in the presence of the IMiD class of small molecules^45^. For example, targeting GSPT1 for degradation by CC-90009 (eragidomide) perturbs the global proteome, activates the integrated stress response (ISR) and induces apoptosis in acute leukemia (AML) cells^46,47^. However, the observation that the VHL-based PROTACs could also induce TNKS1 degradation (Figure S2C) and the addition of IWR1 readily abolished the ability of IWR1-POMA to degrade TNKS (Figure S3C) suggest that IWR1-POMA did not deplete TNKS through affecting translation termination globally. Several lines of evidence further support that the observed growth inhibition by IWR1-POMA is a result of on-target degradation of TNKS. First, HCT116 cells harboring a mutation in one β-catenin allele that cannot be phosphorylated by the DC showed remarkable resistance to IWR1-POMA (Figure 6B and S10A). Second, IWR1-POMA had no effect on the level of cytosolic β-catenin in 293T-TNKS1/2-DKO cells that lack both TNKS1 and TNKS2 (Figure S10B), indicating that IWR1-POMA downregulated β-catenin in a TNKS-dependent manner. Third, the GSPT1/2 degrader CC-90009 had no effect on TNKS and the cytosolic β-catenin levels in DLD-1 cells (Figure S10C). Fourth, we have further exposed DLD-1 cells to CC-90009 and established resistant cells (DLD-1R) that lacked the expression of GSPT1/2 (Figure S10D). TNKS in DLD-1R remained responsive to IWR1 and IWR1-POMA treatments (Figure S10E), and IWR1-POMA was still able to prevent the proliferation of DLD-1R cells effectively (Figure 6B and S10F). As such, IWR1-POMA suppressed WNT-dependent cancer cell proliferation through degrading TNKS but not GSPT1/2. However, consistent with catalytic inhibition or knocking down both TNKS1 and TNKS2 stabilized PTEN and suppressed the growth of RKO cells^48^, IWR1-POMA also suppressed the proliferation of RKO cells that carry full-length APC (Figure 6B and S10G). This result indicates that targeting TNKS by IWR1-POMA can also suppress cancer cell growth through the PTEN-AKT axis independently of the WNT/β-catenin pathway.

To investigate the anticancer potential of IWR1-POMA further, we used DLD-1 and SW480 cells to establish 3D spheroids that recapitulated the cell-cell interactions and the hypoxia core of tumors. Consistent with colony formation assays, IWR1 had no effect on the size and the morphology of these spheroids, while IWR1-POMA suppressed their growth effectively (Figure 6C and S11A). Notably, the outer layer of the spheroids treated by IWR1-POMA had a loose structure incapable of supporting a spherical shape. Instead, they adapted a rather flat architecture with a small core of aggregated cells. Immunostaining showed that the outer layer of these IWR1-POMA-treated spheroids did not contain living cells (Figure 6D). The presence of significant DNA breaks in this region (Figure S11B) is consistent with the observation that APC restoration induced apoptosis in CRC cells^49^.

We have also evaluated the anticancer effects of IWR1-POMA in a human CRC organoid model that preserved the multicellular identity of the tumor more faithfully than immortalized cancer cells. The PDM-7 patient-derived primary CRC cells harbor truncating mutations in APC, making it susceptible to WNT inhibition. We first confirmed that the PDM-7 organoids maintained the heterogeneous nature of the proliferation and the WNT activity within the organoids (Figure S11C). We next confirmed that IWR1-POMA could prevent the formation of PDM-7 organoids whereas IWR1 had little or no effect (Figure S11D). We then treated established PDM-7 organoids with IWR1 or IWR1-POMA and found that, in contrast to IWR1 that did not affect their growth, IWR1-POMA inhibited the proliferation of these organoids with an apoptotic phenotype (Figure 6E and S11E and F) and promoted β-catenin degradation more effectively than IWR1 (Figure 6D). Collectively, removing TNKS scaffolding is important for achieving anticancer efficacy in CRC by targeting TNKS.

## Discussion

In the existing model of WNT/β-catenin signaling, AXIN is a key component of the DC. It interacts with APC to work as a scaffolding protein for the DC to catalyze β-catenin phosphorylation and ubiquitination. In the cytoplasm, TNKS also associates with the DC and controls the AXIN level through PARdU^12–14^. As such, AXIN accumulates upon TNKS inhibition, leading to enhanced β-catenin degradation and reduced WNT/β-catenin pathway activities^12,15^. Traditionally, the formation of AXIN puncta is viewed as a hallmark of functional DC induction^16–18^, wherein the increased local concentration of DC components enhances its “effective activity”^50,51^. Because TNKS inhibition induces large AXIN puncta, it is generally believed that TNKSi supports the formation of highly functional DCs to accelerate the turnover of β-catenin. However, given the newly discovered role of TNKS scaffolding^19,35^, the mechanism by which TNKS controls WNT signaling needs refinement. In this study, we show that AXIN puncta formation relies on the scaffolding but not the catalytic function of TNKS, which is consistent with the previous report that knocking down both TNKS1 and TNKS2 prevented G007-LK to induce AXIN puncta in SW480 cells^18^. However, the function role of TNKS scaffolding in the DC remained unaddressed. Interestingly, blocking proteasomal degradation by MG132 promoted TNKS accumulation but the resulting DC exhibited a different phenotype as compared to that induced by G007-LK^18^. It is likely that the PAR chains helped suppress TNKS aggregation as observed with PARP1.

The DC exhibits characteristics of biomolecular condensates^9,10^. It containing tens to hundreds of AXIN and APC molecules in a ∼1:1 ratio to catalyze β-catenin degradation^50^. Both AXIN and APC contain extensive intrinsically disordered regions, which are likely important for providing a flexible structure to support catalysis. TNKS interacts with AXIN and APC through multivalent binding in the DC. Based on this work, the catalysis-independent function of TNKS in WNT/β-catenin signaling originates from its ability to promote maturation^22,23^ of the DC condensates. Specifically, because TNKS aggregates and forms filaments at high concentrations^34,39,40^, large AXIN puncta containing excess TNKS are rigid and exhibit reduced catalytic activity. In contrast, in the absence of TNKS, the small AXIN complexes are dynamic and catalyze β-catenin degradation more effectively. As such, the size of the AXIN complex does not inform the activity of the DC. This notion is consistent with the previous observation that AXIN2 can also support β-catenin degradation despite having a reduced ability to form puncta^52^. Collectively, our study suggests that TNKS controls WNT/β-catenin signaling through two independent mechanisms: it regulates AXIN homeostasis through catalytic PARylation^12^; meanwhile it dictates the material properties of the DC through molecular scaffolding. The mechanism by which TNKS antagonizes the DC through molecular scaffolding is also reminiscent of the effect of disheveled (DVL) polymerization on the WNT/β-catenin signalosome^53^. However, characterization of the physical properties of reconstituted DC^54^ with TNKS is needed to better understand the nature of these condensates.

The development of TNKSi as an anticancer drug has been discouraged by the lack of significant efficacy in various in vitro and in vivo models. Our data suggest that TNKSd can overcome this limitation by stabilizing AXIN without affecting the DC dynamics. Therefore, TNKSd can provide better suppression of the WNT/β-catenin-controlled genes important for cancer cell proliferation than TNKSi. In particular, we identify several WNT target genes that are suppressed by TNKSd but not TNKSi. Among them, Aurora A is an oncogene frequently amplified in CRC^55^. Because Aurora A binds to and stabilizes c-MYC that is also dysregulated in many cancers^56^, the ability of IWR1-POMA to downregulate both c-MYC and Aurora A more effectively than IWR1 likely contributed to its improved anti-proliferative activity. Another WNT/β-catenin downstream target specifically regulated by TNKSd is cyclin D1. The CDK4/cyclin D1 complex is a key regulator of cell cycle, and CDK4 blockade has been explored as a potential strategy to enhance the anticancer efficacy of TNKSi^57,58^. Interestingly, TNKSd, but not TNKSi, also downregulated CDK4 expression potentially through c-MYC. We speculate that the coordinated action of IWR1-POMA on CDK4 and cyclin D1 additionally contributed to its improved anticancer efficacy. Finally, TNKSi can sensitize cancer cells toward EGFR or MEK inhibition^59,60^. As a more effective suppressor of Wnt/β-catenin signaling, IWR1-POMA may provide a better therapeutic window for treating cancers driven by WNT/β-catenin signaling using a corresponding combinatorial approach.

Some previous studies showed that TNKSi induced reversible intestinal toxicity at high doses without significant antitumor effects in mice^28,61^. However, others found that suppressing the proliferation of *Lgr5*^+^ intestinal stem cells by TNKSi did not affect the morphology of the tissue or cause weight loss^62–64^, and the selective TNKS inhibitor basroparib was well tolerated in a recently disclosed Phase 1 clinical study^65^. As constitutive silencing of TNKS in *APC*-null mice prevented tumorigenesis without damaging the intestines^8^, the observed on-target toxicity of TNKSi may associate with TNKS accumulation at high doses, akin to the cytotoxicity induced by PARP1-trapping upon catalytic inhibition, and reverse allostery^66^ in TNKS has been proposed^40^. Studying TNKS complexes in normal tissues under these conditions may help delineate the source of intestinal toxicity. Finally, targeting the TNKS ARC or SAM domains offers an alternative approach to suppress WNT/β-catenin signaling without promoting TNKS oligomerization^67,68^. Targeting the ARC may further differentiate the WNT-dependent and independent activities of TNKS and provide finer resolution in TNKS inhibition. We are thus cautiously optimistic for continued investment in aggregation-free TNKS-targeting in CRC.

## Materials and Methods

### Cell lines

HAP1, DLD1, HEK293, 239T, HeLa, HT-29, SW480, RKO, and L-Wnt3A cells obtained from Horizon Discovery or ATCC were cultured according to the vendors’ recommended procedures. L-Wnt3A cells were used to generate conditioned medium following the protocol described by Nusse (https://wnt.stanford.edu/purification%23old#medium). PDM-7 obtained from ATCC were grown into organoid according to the procedures provided by ATCC using the Organoid Growth Kit 1A and cell basement membrane for organoid culture from ATCC.

### CRISPR engineering of HAP1 cells

HAP1 cells were transiently transfected with pCRISPaint-NanoLuc-PuroR and pCAS9-mCherry-Frame+2 plasmids and a TNKS1 sgRNA plasmid targeting 5’-TCACTAGGTCTTCTGCTCTGCGG-3’ and selected by puromycin treatment. Cells were then cultured in 96 well plates to generate single colonies and sequenced to select for correct incorporation of nanoluciferase at the C-terminus of TNKS1.

### DLD-1R cells

DLD-1 cells were cultivated with an increasing concentration of CC-90009 (0.1, 1 and 10 µM) to confer resistance to GSPT inhibition. Western analysis confirmed the loss of GSPT1 expression in these cells. GSPT2 is a low-abundance protein in DLD-1 and DLD-1R not detectable by Western blot, which is consistent with a reported quantitative proteomic analysis of DLD-1 cells (GSPT1: 102,567 ppb, GSPT2: 2,495 ppb; https://www.ebi.ac.uk/gxa/experiments/E-PROT-18/Results)^69^.

### Determination of TNKS degradation

HAP1-TNKS1-NanoLuc cells at 30‒50% confluence were treated with different concentrations of PROTAC molecules in triplicates for 16 h. The TNKS1 level was quantified by QUANTI-Luc Gold (Invivogen) with normalization to the total protein content. For high-throughput screens, the cells were seeded into 96-well plates at 2×10^4^ cells per well with 100 μL culture media. The PROTAC molecules dissolved in DMSO were placed in T8+ dispensehead cassettes and then introduced by the TECAN D300e Digital Dispenser with series dilutions under the D300e software control. To determine the DC_50_ value of IWR1-POMA, the dose-response relationship was fitted to a Bayesian Gaussian Processes model in R using code developed by Semenova^70^.

### Immunoblot assays

Cells were washed once with cold PBS and lysed with the SDS lysis buffer (1% SDS, 10 mM HEPES, pH 7.0, 2 mM MgCl_2_ and 500 U universal nuclease). Subcellular protein fractions were obtained by using the Mem-PER Plus Kit (Thermo Fisher Scientific). Briefly, the cell pellets were suspended in the cell permeabilization buffer with cOmplete EDTA-free protease inhibitor cocktail (Sigma) and incubated for 15 min at 4 °C followed by centrifugation at 16,000 × *g* for 15 min to provide the cytosolic fraction. The pellets were then suspended in the cell permeabilization buffer with 0.5% NP-40 and protease inhibitors, passed through a 27½-gauge needle for 30 times, and centrifuge at 16,000 × *g* for 15 min to give the cytosolic fraction, and the pellets were suspended in RIPA buffer with protease inhibitors with sonication to provide the nuclear fraction. Alternatively, cells were incubated in eight volumes of hypotonic buffer (20 mM HEPES-KOH, pH 7.9; 10 mM KCl; 2 mM MgCl₂; 0.3% NP-40; 1 mM DTT; 1 mM EDTA; and 1× protease inhibitor cocktail) on ice for 20 min and then centrifuged at 3,000 × *g* for 10 min at 4°C to provide the cytosolic extracts. Protein concentrations were determined by the BCA assay kit (Thermo Fisher Scientific). A total of 20 μg of proteins were loaded onto the SDS–PAGE gel and then transferred to a nitrocellulose membrane. Nitrocellulose membranes were then blocked with Tris buffered saline containing 0.1% Tween-20 and 5% milk (Bio-Rad). Membranes were incubated with the primary antibodies overnight at 4 °C and then the secondary antibodies for 1 h at room temperature. The blots were developed with or without enhanced chemiluminescence and were exposed on autoradiograph films or imaged by a BioRad Molecular Imager ChemiDoc XRS System.

### Quantitative mass spectrometry

DLD-1 cells at 40% confluence were treated with the drug for 24 h, washed with cold PBS and lysed with the SDS lysis buffer. Protein concentrations were determined by the BCA assay kit (Thermo Fisher Scientific). Two biological replicate samples were prepared with 500 μg of proteins from each sample, reduced with 2 mM DTT for 10 min and alkylated with 50 mM iodoacetamide for 30 min in dark, and then extracted using methanol-chloroform precipitation. The protein pellets were dissolved in 400 μL 8 M urea buffer (8 M urea, 50 mM Tris-HCl, 10 mM EDTA, pH 7.5) and digested by Lys-C (Wako, at a 1:100 enzyme/protein ratio) for 2 h. The urea concentration was then reduced to 2 M using freshly made 100 mM ammonium bicarbonate solution. Proteins were subsequently digested with trypsin (Thermo Fisher Scientific, at 1:100 enzyme/protein ratio) overnight. Peptides were desalted with Oasis HLB cartridges (Waters) and re-suspended in 200 mM HEPES (pH 8.5) to a final concentration of 1 μg/μL. For each sample, 100 μg of peptides were reacted with the corresponding amine-based TMT reagents (Thermo Fisher Scientific) for 1 h. The reactions were quenched with 5% hydroxylamine solution and were combined. Samples were then desalted and a reverse-phase fractionation spin column (Pierce) was used according to the manufacturer’s directions to fractionate the sample into 8 fractions. The fractions were dried in a SpeedVac and reconstituted in a 2% acetonitrile, 0.1% TFA buffer followed by injecting onto an Orbitrap Fusion Lumos mass spectrometer coupled to an Ultimate 3000 RSLC-Nano liquid chromatography system. Samples were injected onto a 75 μm×75-cm EasySpray column (Thermo) and eluted with a gradient from 0→28% buffer B over 180 min. Buffer A contained 2% (v/v) acetonitrile and 0.1% formic acid in water, and buffer B contained 80% (v/v) acetonitrile, 10% (v/v) trifluoroethanol, and 0.1% formic acid in water. The mass spectrometer operated in positive ion mode with a source voltage of 1.8 kV and an ion transfer tube temperature of 275 °C. MS scans were acquired at 120,000 resolution in the Orbitrap and top speed mode was used for SPS-MS3 analysis with a cycle time of 2.5 s. MS2 was performed with CID with a collision energy of 35%. The top 10 fragments were selected for MS3 fragmentation using HCD, with a collision energy of 55%. Dynamic exclusion was set for 25 s after an ion was selected for fragmentation. The raw MS data files were analyzed using Proteome Discoverer v2.4 (Thermo), with peptide identification performed using Sequest HT searching against the human protein database from UniProt. Fragment and precursor tolerances of 10 ppm and 0.6 Da were specified, and three missed cleavages were allowed. Carbamidomethylation of Cys and TMT10plex labelling of N-termini and Lys side-chains were set as fixed modifications, with oxidation of Met set as a variable modification. Protein abundances were determined based on the sum of the signal-to-noise ratios of the reporter ions for all peptides matched to each protein. The false-discovery rate (FDR) cutoff was 1% for all peptides. Statistical analyses (one-way ANOVA and two-sided unpaired t-tests) were performed in R using the peptide data. The MS data have been deposited in the MassIVE repository with the dataset identifier MSV000089098.

### Luciferase reporter assays

L-Wnt3A-STF cells, or HeLa, DLD-1, or SW480 cells expressing STF and Renilla-luciferase at 30‒50% confluence were treated with the indicated compound for 16 h. The STF-firefly and Renilla luciferase activities were measured by the Dual Luciferase kit (Promega). The WNT/β-catenin pathway activities were determined by normalizing the STF-firefly activity to the Renilla activity or the total protein level. Briefly, cells were transfected with or without STF plasmid (Addgene) and pRL Renilla luciferase plasmid (Addgene) using Lipofectamine 3000 (Thermo Fisher Scientific) in Opti-MEM for 5 h and empty cloning vector as negative control. After incubating in cell growth media for 3 h, cells were collected and reseeded into 24 or 96-well plates. After incubating for 4 h, cells were treated with the indicated small molecules for 16–24 h and then washed with PBS, and then lysed with Passive Lysis Buffer (Promega) before luciferase activity was measured according to the manufacturer’s instructions. Statistical analyses (one-way ANOVA and two-sided unpaired t-tests) were performed in GraphPad Prism.

### BRET analysis

HAP1-TNKS1-NanoLuc cells (2×10^4^ cells) treated with MG132 (1 μM) for 2 h or 293T cells (1×10^4^ cells) transfected with TNKS1-NanoLuc plasmid (25 ng) for 24 h were treated with different concentrations of IWR1 in the presence of IWR1-POMA (0.5 μM) or IWR1-PEG3-NCT (0.5 μM) for 2 h in Opti-MEM without phenol red in 96-well plates. NanoBRET NanoGlo Substrate (Promega) was then added and the fluorescence signals were measured at 450 and 610 nm on a BioTek Synergy Neo2 microplate reader. The BRET signals were determined as the 610 nm (acceptor) emission over 450 nm (donor) emission ratio after background correction.

### Immunofluorescence microscopy

HeLa cells at 30‒50% confluence in a 35 mm dish were transfected with 30 ng of the indicated plasmids for 6 h and then treated with DMSO, IWR (3 μM) or IWR1-POMA (3 μM) for 16 h. Images were obtained using a Zeiss LSM 880 Airyscan confocal laser scanning microscope. FRAP analysis was performed at room temperature in the DMEM media without phenol red and with HEPES supplement and the indicated drugs. Defined regions were photobleached at a specific wavelength using the 405 nm or 561 nm laser, and the fluorescence intensities in these regions were collected every 2 s and normalized to the initial intensity before bleaching. The fluorescent signal intensities were determined in ImageJ and analyzed in GraphPad Prism.

### 2D colony formation assays

DLD-1, SW480, HT-29, HCT116, RKO, or DLD-1R cells were seeded into 6-well plates at 500‒2000 cells/well with 10% FBS unless otherwise mentioned in the growth medium. Compounds were added at the indicated concentrations 16 h after seeding, and the growth medium was replenished every 3 d until colony formation was observed. The colonies were fixed with 4% formalin in PBS and stained by a solution of 0.5% crystal violet in 50% methanol solution.

### 3D spheroid formation assay

DLD-1 or SW480 cells were seeded at 1,000 cells/well into 96-well plates with 10% FBS in the Hyclone X growth medium. For DLD-1 cells, 3D spheroids formed 1 d after seeding. For SW480 cells, rat tail collagen I (Gibco) was added to provide the extracellular matrix, and 3D spheroids formed 5 d after seeding. Compounds were then added at the indicated concentrations. Medium with the indicated drug was replenished every 3 d, and the size of the spheroids were measured at the indicated time by imaging on a Cytation 5 Cell Imaging Multimode Reader and analyzed by Image J. The spheroids at the endpoints were fix with 4% formalin in PBS, sectioned and stained with the indicated markers after parafilm embedding.

### CRC organoid formation assay

The organoids were cultured according to the procedures detailed by ATCC. Briefly, dissociated PDM-7 single cells or organoid fragments of about 200 μM diameter in size were embedded at 5,000,000 cells/mL in the cell base membrane and dispensed as small droplets onto warm 6-well plates. After solidification, the domes were covered with the advanced DMEM:F12 supplemented with HEPES, L-glutamine and B-27, noggin, gastrin, N-acetyl-cysteine, EGF, nicotinamide, A 83-01, and SB 202190 (ATCC formulation 1). The ROCK inhibitor Y-27632 (10 μM) was also included for the first 3 d of subculture. The organoids were treated with the drug 3 days after seeding for 5 days in the single-cell model and 1 day after seeding for 5 days in the organoid fragment model.

### Synthetic procedures

#### Synthesis of IWR1-TP4-Poma (IWR1-POMA)

To a solution of [*1127442-97-0*]^15^ (351 mg, 0.8 mmol, 1.0 equiv) in methylene chloride (3 mL) was added methanesulfonyl chloride (183 mg, 1.6 mmol, 2.0 equiv), triethylamine (0.33 mL, 2.4 mmol, 3.0 equiv) at 0 °C. After stirring for 2 h, the reaction mixture was concentrated, the residual was re-dissolved in *N*,*N*-dimethylformamide (3 mL), and sodium azide (208 mg, 3.2 mmol, 4.0 equiv) was then added. After stirring at 50 °C overnight, the reaction was quenched with water and extracted with ethyl acetate for three times. The combined organic layers were washed with brine, dried over sodium sulfate, concentrated, and purified by silica gel flash column chromatography to give azido-IWR2 (335 mg, 90% yield) as a white powder. ^1^H NMR (400 MHz, CDCl_3_) δ 10.74 (s, 1H), 8.90 (dd, *J* = 6.0, 3.0 Hz, 1H), 8.78 (d, *J* = 4.4 Hz, 1H), 8.09 (d, *J* = 8.1 Hz, 2H), 7.63–7.56 (m, 2H), 7.47 (d, *J* = 4.4 Hz, 1H), 7.36 (d, *J* = 8.2 Hz, 2H), 6.28 (t, *J* = 1.8 Hz, 2H), 4.82 (s, 2H), 3.54–3.50 (m, 2H), 3.46 (dd, *J* = 3.1, 1.6 Hz, 2H), 1.79 (d, *J* = 8.6 Hz, 1H), 1.62 (d, *J* = 8.8 Hz, 1H); MS (ESI) calcd for C_26_H_21_N_6_O_3_ (M+H)^+^ 465.2, found 465.2.

To a solution of azido-IWR2 (4.2 mg, 0.009 mmol, 1.0 equiv) in dimethyl sulfoxide was added [*2138439-58-2*]^71^ (5 mg, 0.01 mmol, 1.1 equiv), copper(II) sulfate pentahydrate (4.5 mg, 0.018 mmol, 2.0 equiv) and (+)-sodium l-ascorbate (7.2 mg, 0.036 mmol, 4.0 equiv). After stirring at 80 °C overnight, the reaction was quenched with saturated ammonium chloride and extracted with methylene chloride for three times. The combined organic layers were washed with brine, dried over sodium sulfate, concentrated, and purified by silica gel flash column chromatography followed by preparative HPLC to give IWR1-TP4-Poma (IWR1-POMA) (5.4 mg, 63% yield) as a yellow solid. ^1^H NMR (400 MHz, CDCl_3_) δ 10.75 (s, 1H), 8.94 (dd, *J* = 7.0, 1.9 Hz, 1H), 8.77 (d, *J* = 4.4 Hz, 1H), 8.38 (s, 1H), 8.14–8.07 (m, 2H), 7.73–7.58 (m, 3H), 7.44 (dd, *J* = 8.5, 7.1 Hz, 1H), 7.40–7.34 (m, 2H), 7.07 (dd, *J* = 7.9, 5.7 Hz, 2H), 6.85 (d, *J* = 8.5 Hz, 1H), 6.29 (t, *J* = 1.9 Hz, 2H), 6.02 (s, 2H), 4.87 (dd, *J* = 12.0, 5.4 Hz, 1H), 4.70 (s, 2H), 3.74–3.58 (m, 14H), 3.54 (dq, *J* = 3.5, 1.7 Hz, 2H), 3.49 (dd, *J* = 3.0, 1.6 Hz, 2H), 3.41 (t, *J* = 5.3 Hz, 2H), 2.89–2.66 (m, 3H), 2.14–2.04 (m, 1H), 1.82 (dt, *J* = 8.9, 1.7 Hz, 1H), 1.65 (d, *J* = 8.9 Hz, 1H); MS (ESI) calcd for C_50_H_50_N_9_O_11_ (M+H)^+^ 952.4, found 952.3.

#### Synthesis of IWR1-TP(n)-Poma

Prepared using the same method as described for IWR1-TP4-Poma using pomalidomide derivatives with different PEG chain length.

##### IWR1-TP1-Poma

7.4 mg, 41% yield. ^1^H NMR (400 MHz, CDCl_3_) δ 10.73 (s, 1H), 8.95 (dd, *J* = 6.3, 2.7 Hz, 1H), 8.74 (d, *J* = 4.3 Hz, 1H), 8.12 (dt, *J* = 9.1, 1.8 Hz, 3H), 7.71–7.61 (m, 3H), 7.44 (ddd, *J* = 8.4, 7.0, 1.1 Hz, 1H), 7.40–7.36 (m, 2H), 7.07 (d, *J* = 7.1 Hz, 1H), 7.01 (d, *J* = 4.5 Hz, 1H), 6.86 (d, *J* = 8.5 Hz, 1H), 6.29 (q, *J* = 1.6 Hz, 2H), 6.03 (s, 2H), 4.84 (dd, *J* = 12.0, 5.4 Hz, 1H), 4.71 (s, 2H), 3.74 (d, *J* = 5.3 Hz, 2H), 3.54 (dq, *J* = 3.3, 1.6 Hz, 2H), 3.50 (d, *J* = 1.1 Hz, 2H), 3.46 (d, *J* = 6.4 Hz, 2H), 2.89–2.59 (m, 3H), 1.82 (dt, *J* = 8.9, 1.5 Hz, 1H), 1.64 (d, *J* = 8.9 Hz, 1H); MS (ESI) calcd for C_44_H_38_N_9_O_8_ (M+H)^+^ 820.3, found 820.2.

##### IWR1-TP2-Poma

10.2 mg, 76% yield. ^1^H NMR (400 MHz, CDCl_3_) δ 10.71 (s, 1H), 8.92 (dd, *J* = 7.4, 1.6 Hz, 1H), 8.72 (d, *J* = 4.4 Hz, 1H), 8.25 (s, 1H), 8.14–8.07 (m, 2H), 7.69–7.57 (m, 3H), 7.43–7.34 (m, 3H), 7.03 (dd, *J* = 5.9, 3.7 Hz, 2H), 6.81 (d, *J* = 8.5 Hz, 1H), 6.29 (t, *J* = 1.9 Hz, 2H), 6.01 (s, 2H), 4.87 (dd, *J* = 12.0, 5.4 Hz, 1H), 4.71 (s, 2H), 3.70 (dd, *J* = 5.9, 3.2 Hz, 2H), 3.66–3.60 (m, 4H), 3.54 (dq, *J* = 3.5, 1.7 Hz, 2H), 3.49 (d, *J* = 1.6 Hz, 4H), 3.35 (t, *J* = 5.3 Hz, 2H), 2.97–2.64 (m, 3H), 1.82 (dt, *J* = 8.9, 1.7 Hz, 1H), 1.64 (d, *J* = 8.7 Hz, 1H); MS (ESI) calcd for C_46_H_42_N_9_O_9_ (M+H)^+^ 864.3, found 864.3. *IWR1-TP3-Poma:* 11.0 mg, 95 % yield. ^1^H NMR (400 MHz, CDCl_3_) δ 10.74 (s, 1H), 8.93 (dd, *J* = 6.1, 2.8 Hz, 1H), 8.78 (d, *J* = 4.4 Hz, 1H), 8.55 (s, 1H), 8.15–8.06 (m, 2H), 7.70–7.58 (m, 3H), 7.44 (dd, *J* = 8.5, 7.1 Hz, 1H), 7.40–7.35 (m, 2H), 7.09 (d, *J* = 4.4 Hz, 1H), 7.05 (d, *J* = 7.1 Hz, 1H), 6.84 (d, *J* = 8.5 Hz, 1H), 6.29 (t, *J* = 1.9 Hz, 2H), 6.01 (s, 2H), 4.93–4.85 (m, 1H), 4.70 (s, 2H), 3.76–3.59 (m, 10H), 3.54 (dq, *J* = 3.4, 1.6 Hz, 2H), 3.49 (dd, *J* = 3.0, 1.6 Hz, 2H), 3.39 (t, *J* = 5.2 Hz, 2H), 2.95–2.66 (m, 3H), 2.10 (td, *J* = 9.8, 9.0, 3.3 Hz, 1H), 1.82 (dt, *J* = 8.9, 1.7 Hz, 1H), 1.65 (d, *J* = 8.7 Hz, 1H); MS (ESI) calcd for C_48_H_46_N_9_O_10_ (M+H)^+^ 908.3, found 908.3.

##### IWR1-TP5-Poma

7.6 mg, 78% yield. ^1^H NMR (400 MHz, CDCl_3_) δ 10.76 (s, 1H), 8.95 (dd, *J* = 7.5, 1.4 Hz, 1H), 8.77 (d, *J* = 4.4 Hz, 1H), 8.51 (s, 1H), 8.15–8.06 (m, 2H), 7.77–7.62 (m, 3H), 7.46 (dd, *J* = 8.5, 7.1 Hz, 1H), 7.42–7.33 (m, 2H), 7.21–7.14 (m, 2H), 7.10–7.03 (m, 2H), 6.87 (d, *J* = 8.5 Hz, 1H), 6.29 (t, *J* = 1.8 Hz, 2H), 6.03 (s, 2H), 4.88 (dd, *J* = 11.9, 5.4 Hz, 1H), 4.68 (s, 2H), 3.75–3.58 (m, 18H), 3.54 (dq, *J* = 3.4, 1.7 Hz, 2H), 3.48 (dd, *J* = 3.0, 1.6 Hz, 2H), 3.41 (t, *J* = 5.2 Hz, 2H), 2.91–2.63 (m, 3H), 2.10 (dt, *J* = 10.5, 4.1 Hz, 1H), 1.82 (dt, *J* = 8.9, 1.7 Hz, 1H), 1.65 (dd, *J* = 8.8, 1.6 Hz, 1H); MS (ESI) calcd for C_52_H_54_N_9_O_12_ (M+H)^+^ 996.4, found 996.4.

##### IWR1-TP6-Poma

13.1 mg, 91% yield. ^1^H NMR (400 MHz, CDCl_3_) δ 10.75 (s, 1H), 8.94 (d, *J* = 7.5 Hz, 1H), 8.77 (d, *J* = 4.3 Hz, 1H), 8.61 (s, 1H), 8.11 (d, *J* = 8.4 Hz, 2H), 7.84–7.60 (m, 3H), 7.44 (dd, *J* = 8.5, 7.1 Hz, 1H), 7.37 (d, *J* = 8.3 Hz, 2H), 7.06 (dd, *J* = 9.4, 5.7 Hz, 2H), 6.86 (d, *J* = 8.5 Hz, 1H), 6.29 (t, *J* = 1.9 Hz, 2H), 6.04 (s, 2H), 4.88 (dd, *J* = 11.7, 5.4 Hz, 1H), 4.69 (s, 2H), 3.68 (t, *J* = 5.1 Hz, 4H), 3.64–3.56 (m, 20H), 3.55–3.51 (m, 2H), 3.48 (d, *J* = 2.1 Hz, 2H), 3.41 (t, *J* = 5.3 Hz, 2H), 2.89–2.69 (m, 3H), 2.14– 2.04 (m, 1H), 1.85–1.77 (m, 1H), 1.64 (d, *J* = 8.9 Hz, 1H); MS (ESI) calcd for C_54_H_58_N_9_O_13_ (M+H)^+^ 1040.4, found 1040.4.

#### Synthesis of IWR1-P(n)-Poma

To a solution of [*2472645-01-3*]^72^ (1.0 equiv) in methylene chloride was added amino-PEG(n)-pomalidomide (1.05 equiv) followed by 4Å molecular sieves. After stirring at 23 °C overnight, sodium triacetoxyborohydride (20 equiv) was added and the reaction was stirred for 4 h before quenched with water. The mixture was extracted with methylene chloride for three times. The combined organic layers were washed with brine, dried over sodium sulfate, concentrated, and purified by silica gel flash column chromatography followed by preparative HPLC to give IWR1-P(n)-Poma as a yellow solid. *IWR1-P1-Poma:* 16.0 mg, 78% yield. ^1^H NMR (500 MHz, CDCl_3_) δ 10.48 (s, 1H), 9.09 (s, 1H), 8.67 (t, *J* = 6.1 Hz, 2H), 7.97 (d, *J* = 8.1 Hz, 2H), 7.55 (d, *J* = 8.5 Hz, 2H), 7.43 (t, *J* = 8.0 Hz, 1H), 7.36 (t, *J* = 7.7 Hz, 1H), 7.32 (d, *J* = 8.1 Hz, 2H), 6.96 (d, *J* = 7.1 Hz, 1H), 6.76 (d, *J* = 8.6 Hz, 1H), 6.27 (t, *J* = 1.9 Hz, 2H), 4.74 (dd, *J* = 13.1, 5.3 Hz, 1H), 4.59 (s, 2H), 3.74 (s, 2H), 3.62 (t, *J* = 5.0 Hz, 2H), 3.54–3.49 (m, 2H), 3.47 (s, 4H), 3.34 (d, *J* = 5.6 Hz, 2H), 3.28–3.19 (m, 2H), 2.79–2.38 (m, 3H), 1.93 (d, *J* = 12.4 Hz, 1H), 1.80 (dt, *J* = 8.9, 1.7 Hz, 1H), 1.63 (d, *J* = 8.8 Hz, 1H); MS (ESI) calcd for C_43_H_40_N_7_O_8_ (M+H)^+^ 782.3, found 782.2.

##### IWR1-P2-Poma

13.1 mg, 56% yield. ^1^H NMR (500 MHz, CDCl_3_) δ 10.53 (s, 1H), 9.39–9.14 (m, 1H), 8.71 (dd, *J* = 23.5, 5.9 Hz, 2H), 8.12–7.91 (m, 2H), 7.60 (t, *J* = 7.4 Hz, 2H), 7.49 (t, *J* = 8.2 Hz, 1H), 7.36 (d, *J* = 8.0 Hz, 2H), 7.32–7.23 (m, 2H), 7.16 (dd, *J* = 12.7, 7.1 Hz, 1H), 6.95 (d, *J* = 7.0 Hz, 1H), 6.60 (d, *J* = 8.5 Hz, 1H), 6.29 (t, *J* = 1.9 Hz, 2H), 4.91 (dd, *J* = 12.7, 5.4 Hz, 1H), 4.69 (s, 2H), 3.85 (t, *J* = 4.8 Hz, 2H), 3.68 (d, *J* = 4.3 Hz, 3H), 3.63 (dt, *J* = 10.2, 4.1 Hz, 4H), 3.58–3.45 (m, 6H), 3.39 (s, 2H), 3.22 (t, *J* = 5.1 Hz, 2H), 2.85–2.67 (m, 2H), 2.62 (td, *J* = 12.8, 4.7 Hz, 1H), 2.08–1.96 (m, 1H), 1.82 (dt, *J* = 9.0, 1.8 Hz, 1H), 1.65 (d, *J* = 8.8 Hz, 1H); MS (ESI) calcd for C_45_H_44_N_7_O_9_ (M+H)^+^ 826.3, found 826.3.

##### IWR1-P3-Poma

13.2 mg, 72% yield. ^1^H NMR (500 MHz, CDCl_3_) δ 10.65 (s, 1H), 9.23 (s, 1H), 8.83 (d, *J* = 7.6 Hz, 1H), 8.75 (d, *J* = 4.0 Hz, 1H), 8.10–8.01 (m, 2H), 7.66 (t, *J* = 7.3 Hz, 2H), 7.56 (t, *J* = 8.2 Hz, 1H), 7.45–7.31 (m, 3H), 7.00 (d, *J* = 7.1 Hz, 1H), 6.74 (d, *J* = 8.6 Hz, 1H), 6.29 (d, *J* = 1.8 Hz, 2H), 4.92–4.82 (m, 1H), 4.69 (s, 2H), 3.79 (s, 2H), 3.67–3.51 (m, 14H), 3.51–3.43 (m, 4H), 3.28 (s, 2H), 2.89–2.59 (m, 3H), 2.05 (q, *J* = 9.1, 7.1 Hz, 1H), 1.85–1.77 (m, 1H), 1.64 (d, *J* = 8.9 Hz, 1H); MS (ESI) calcd for C_47_H_48_N_7_O_10_ (M+H)^+^ 870.3, found 870.3.

##### IWR1-P4-Poma

15.6 mg, 68% yield. ^1^H NMR (500 MHz, CDCl_3_) δ 10.66 (s, 1H), 9.02 (s, 1H), 8.85 (d, *J* = 7.7 Hz, 1H), 8.76 (d, *J* = 3.9 Hz, 1H), 8.07 (d, *J* = 8.2 Hz, 2H), 7.77–7.65 (m, 2H), 7.56 (t, *J* = 8.1 Hz, 1H), 7.36 (d, *J* = 8.0 Hz, 2H), 7.32 (t, *J* = 7.8 Hz, 1H), 6.97 (d, *J* = 7.0 Hz, 1H), 6.64 (d, *J* = 8.5 Hz, 1H), 6.29 (d, *J* = 2.0 Hz, 2H), 4.87 (q, *J* = 5.7 Hz, 1H), 4.71 (s, 2H), 3.80 (s, 2H), 3.68–3.40 (m, 22H), 3.32 (s, 2H), 3.11 (d, *J* = 5.3 Hz, 2H), 2.88–2.58 (m, 3H), 2.12–2.00 (m, 1H), 1.81 (d, *J* = 8.9 Hz, 1H), 1.64 (d, *J* = 8.8 Hz, 1H); MS (ESI) calcd for C_49_H_52_N_7_O_11_ (M+H)^+^ 914.4, found 914.3.

##### IWR1-P5-Poma

13.6 mg, 53% yield. ^1^H NMR (500 MHz, CDCl_3_) δ 10.72 (s, 1H), 8.89 (d, *J* = 7.7 Hz, 1H), 8.86 (s, 2H), 8.81 (d, *J* = 4.1 Hz, 1H), 8.10 (d, *J* = 8.0 Hz, 2H), 7.74 (dd, *J* = 13.1, 6.4 Hz, 2H), 7.60 (t, *J* = 8.1 Hz, 1H), 7.37 (dd, *J* = 8.4, 3.9 Hz, 3H), 7.24 (d, *J* = 7.6 Hz, 1H), 7.16 (dd, *J* = 12.6, 7.1 Hz, 1H), 7.01 (d, *J* = 7.1 Hz, 1H), 6.74 (d, *J* = 8.5 Hz, 1H), 6.29 (d, *J* = 1.9 Hz, 2H), 4.99–4.83 (m, 1H), 4.75 (s, 2H), 3.80 (s, 2H), 3.70–3.41 (m, 24H), 3.32 (s, 2H), 3.26 (t, *J* = 4.2 Hz, 2H), 2.76 (ddd, *J* = 61.7, 10.7, 4.5 Hz, 3H), 2.08 (dq, *J* = 7.6, 4.1, 2.7 Hz, 1H), 1.81 (d, *J* = 9.0 Hz, 1H), 1.64 (d, *J* = 8.8 Hz, 1H); MS (ESI) calcd for C_51_H_56_N_7_O_12_ (M+H)^+^ 958.4, found 958.4.

##### IWR1-P6-Poma

9.2 mg, 40% yield. ^1^H NMR (500 MHz, CDCl_3_) δ 10.74 (s, 1H), 8.90 (d, *J* = 7.5 Hz, 2H), 8.83 (d, *J* = 4.1 Hz, 1H), 8.11 (d, *J* = 7.9 Hz, 2H), 7.76 (dd, *J* = 11.8, 6.5 Hz, 2H), 7.60 (t, *J* = 8.1 Hz, 1H), 7.38 (dd, *J* = 8.6, 2.2 Hz, 3H), 7.02 (d, *J* = 7.1 Hz, 1H), 6.73 (d, *J* = 8.5 Hz, 1H), 6.29 (t, *J* = 1.8 Hz, 2H), 4.89 (q, *J* = 5.5, 5.1 Hz, 1H), 4.77 (t, *J* = 9.9 Hz, 2H), 3.83 (s, 2H), 3.68–3.44 (m, 28H), 3.35 (s, 2H), 3.28 (d, *J* = 5.1 Hz, 2H), 2.93–2.62 (m, 3H), 2.18–2.01 (m, 1H), 1.85–1.79 (m, 1H), 1.65 (d, *J* = 8.9 Hz, 1H); MS (ESI) calcd for C_53_H_60_N_7_O_13_ (M+H)^+^ 1002.4, found 1002.4.

#### Synthesis of IWR1-P(n)-VHL

Prepared as a white powder using the same method as described for IWR1-P(n)-Poma using amino-PEG(n)-VH032 derivatives^73^ with different PEG chain length.

##### IWR1-P1-VHL

2.4 mg, 9% yield. ^1^H NMR (400 MHz, CDCl_3_) δ 10.75 (s, 1H), 8.97 (d, *J* = 7.7 Hz, 1H), 8.83 (d, *J* = 4.4 Hz, 1H), 8.75 (s, 1H), 8.20–8.00 (m, 2H), 7.89 (d, *J* = 4.5 Hz, 1H), 7.84 (d, *J* = 8.6 Hz, 1H), 7.71 (t, *J* = 8.1 Hz, 1H), 7.41–7.32 (m, 2H), 7.29 (s, 4H), 6.66 (d, *J* = 8.7 Hz, 1H), 6.29 (t, *J* = 1.9 Hz, 2H), 4.92 (s, 2H), 4.68–4.43 (m, 4H), 4.28 (dd, *J* = 15.2, 5.2 Hz, 1H), 4.02 (d, *J* = 11.4 Hz, 1H), 3.87 (s, 1H), 3.76 (s, 3H), 3.59 (d, *J* = 11.0 Hz, 1H), 3.55 (s, 3H), 3.49 (t, *J* = 2.3 Hz, 2H), 3.30 (q, *J* = 18.2, 15.0 Hz, 5H), 2.49 (s, 3H), 1.82 (d, *J* = 8.9 Hz, 1H), 1.65 (d, *J* = 9.0 Hz, 2H), 0.96 (s, 9H); MS (ESI) calcd for C_53_H_59_N_8_O_8_S (M+H)^+^ 967.4, found 967.4.

##### IWR1-P2-VHL

5.3 mg, 18% yield. ^1^H NMR (400 MHz, CDCl_3_) δ 10.76 (t, *J* = 2.8 Hz, 1H), 8.96 (q, *J* = 4.6, 3.7 Hz, 2H), 8.85 (q, *J* = 3.2, 2.3 Hz, 1H), 8.10 (dt, *J* = 8.7, 2.7 Hz, 2H), 7.99–7.58 (m, 5H), 7.42–7.33 (m, 4H), 7.31 (t, *J* = 2.8 Hz, 2H), 7.02 (d, *J* = 8.6 Hz, 1H), 6.47–6.13 (m, 2H), 4.90 (d, *J* = 5.4 Hz, 2H), 4.58 (td, *J* = 19.0, 16.8, 11.0 Hz, 5H), 4.31 (dd, *J* = 12.0, 8.0 Hz, 1H), 3.99 (d, *J* = 11.2 Hz, 1H), 3.85 (s, 3H), 3.70–3.60 (m, 7H), 3.55 (s, 3H), 3.50 (t, *J* = 3.2 Hz, 4H), 3.37–3.23 (m, 3H), 2.61–2.36 (m, 6H), 2.25 (d, *J* = 26.9 Hz, 3H), 1.82 (d, *J* = 9.0 Hz, 1H), 1.70–1.53 (m, 1H), 1.32 (td, *J* = 7.3, 3.4 Hz, 4H), 0.97 (d, *J* = 3.0 Hz, 10H); MS (ESI) calcd for C_55_H_63_N_8_O_9_S (M+H)^+^ 1011.4, found 1011.4.

##### IWR1-P3-VHL

4.7 mg, 23% yield. ^1^H NMR (400 MHz, CDCl_3_) δ 10.76 (s, 1H), 9.15 (s, 1H), 8.96 (d, *J* = 7.5 Hz, 1H), 8.89 (d, *J* = 4.1 Hz, 1H), 8.10 (dd, *J* = 8.6, 2.1 Hz, 2H), 7.95–7.87 (m, 1H), 7.79 (d, *J* = 8.6 Hz, 1H), 7.70 (t, *J* = 8.2 Hz, 1H), 7.38 (d, *J* = 7.9 Hz, 4H), 7.32 (d, *J* = 7.7 Hz, 2H), 6.29 (s, 2H), 4.91 (s, 2H), 4.76–4.39 (m, 4H), 3.86 (s, 2H), 3.80–3.41 (m, 17H), 3.37–3.13 (m, 2H), 2.54 (d, *J* = 2.0 Hz, 3H), 2.43 (s, 2H), 2.27 (s, 2H), 1.82 (d, *J* = 8.9 Hz, 1H), 1.65 (d, *J* = 8.8 Hz, 1H), 1.37–1.29 (m, 3H), 0.96 (s, 9H); MS (ESI) calcd for C_57_H_67_N_8_O_10_S (M+H)^+^ 1055.5, found 1055.4.

##### IWR1-P4-VHL

3.3 mg, 32% yield. ^1^H NMR (400 MHz, CDCl_3_) δ 10.72 (s, 1H), 8.89 (d, *J* = 7.7 Hz, 1H), 8.85–8.77 (m, 2H), 8.02 (d, *J* = 8.2 Hz, 2H), 7.81 (dd, *J* = 20.6, 6.6 Hz, 2H), 7.61 (t, *J* = 8.1 Hz, 1H), 7.36– 7.23 (m, 6H), 6.91 (d, *J* = 10.9 Hz, 1H), 6.21 (t, *J* = 1.9 Hz, 2H), 4.81 (s, 2H), 4.66–4.18 (m, 5H), 3.82 (d, *J* = 20.1 Hz, 3H), 3.65–3.37 (m, 23H), 3.25 (s, 2H), 2.46 (s, 3H), 1.75 (d, *J* = 8.7 Hz, 1H), 1.57 (d, *J* = 8.9 Hz, 1H), 1.29 (t, *J* = 7.0 Hz, 3H), 0.86 (s, 9H); MS (ESI) calcd for C_59_H_71_N_8_O_11_S (M+H)^+^ 1099.5, found 1099.5.

##### IWR1-P5-VHL

6.9 mg, 32% yield. ^1^H NMR (400 MHz, CDCl_3_) δ 10.70 (s, 1H), 8.94 (s, 1H), 8.90 (d, *J* = 7.7 Hz, 1H), 8.82 (d, *J* = 4.0 Hz, 1H), 8.04 (d, *J* = 8.1 Hz, 2H), 7.88–7.85 (m, 1H), 7.76 (d, *J* = 8.6 Hz, 1H), 7.63 (t, *J* = 8.1 Hz, 1H), 7.30 (dd, *J* = 12.7, 7.1 Hz, 7H), 6.22 (d, *J* = 2.2 Hz, 2H), 4.85 (s, 2H), 4.72–4.35 (m, 4H), 4.27 (d, *J* = 15.0 Hz, 1H), 3.82 (s, 2H), 3.72–3.28 (m, 21H), 3.24 (d, *J* = 7.6 Hz, 3H), 2.46 (s, 3H), 1.75 (d, *J* = 8.9 Hz, 1H), 1.58 (d, *J* = 8.9 Hz, 1H), 1.28 (t, *J* = 6.8 Hz, 3H), 0.88 (s, 9H); MS (ESI) calcd for C_61_H_75_N_8_O_12_S (M+H)^+^ 1143.5, found 1143.4.

##### IWR1-P6-VHL

6.7 mg, 20% yield. ^1^H NMR (400 MHz, CDCl_3_) δ 10.71 (s, 1H), 9.02 (s, 1H), 8.90 (d, *J* = 7.7 Hz, 1H), 8.83 (d, *J* = 4.3 Hz, 1H), 8.10–7.99 (m, 2H), 7.86 (d, *J* = 4.4 Hz, 1H), 7.76 (d, *J* = 8.5 Hz, 1H), 7.64 (t, *J* = 8.1 Hz, 1H), 7.38–7.25 (m, 7H), 6.22 (t, *J* = 1.9 Hz, 2H), 4.87 (s, 2H), 4.50 (ddd, *J* = 35.0, 24.8, 16.4 Hz, 4H), 4.27 (dd, *J* = 15.2, 4.9 Hz, 1H), 4.02 (d, *J* = 11.3 Hz, 1H), 3.82 (d, *J* = 5.4 Hz, 2H), 3.68–3.38 (m, 32H), 3.27 (d, *J* = 7.3 Hz, 2H), 2.47 (s, 3H), 1.75 (dt, *J* = 9.0, 1.7 Hz, 1H), 1.58 (d, *J* = 8.9 Hz, 1H), 1.30 (t, *J* = 7.0 Hz, 3H), 0.89 (s, 9H); MS (ESI) calcd for C_63_H_79_N_8_O_13_S (M+H)^+^ 1187.5, found 1187.5.

#### Synthesis of IWR6-P(n)-Poma and IWR6-P(n)-VHL

Prepared using the same methods as described for IWR1-P(n)-Poma and IWR1-P(n)-VHL using [*1801530-64-2*]^74^.

#### Synthesis of IWR1-C6-Poma

Prepared using the same methods as described for IWR1-P(n)-Poma using [*2093386-50-4*]^75^. MS (ESI) calcd for C_45_H_44_N_7_O_7_ (M+H)^+^ 794.3, found 794.3.

#### Synthesis of IWR1-R-Olena

To a solution of [*2472645-01-3*]^72^ (40 mg, 0.09 mmol, 1.0 equiv) in methylene chloride (0.5 mL) was added [*852180-47-3*]^76^ (28 mg, 0.095 mmol, 1.05 equiv) followed by 4Å molecular sieves (400 mg). After stirring at 23 °C overnight, sodium triacetoxyborohydride (48 mg, 0.23 mmol, 2.5 equiv) was added and the reaction was stirred for 4 h before quenched with water. The mixture was extracted with methylene chloride for three times. The combined organic layers were washed with brine, dried over sodium sulfate, concentrated, and purified by silica gel flash column chromatography followed by preparative HPLC to give IWR1-RL (45 mg, 69% yield) as a white powder. ^1^H NMR (400 MHz, CDCl_3_) δ 10.82 (s, 1H), 8.89 (d, *J* = 7.8 Hz, 1H), 8.78 (dd, *J* = 4.4, 1.2 Hz, 1H), 8.16–8.03 (m, 2H), 7.67–7.62 (m, 1H), 7.61–7.53 (m, 2H), 7.41–7.34 (m, 2H), 7.27 (d, *J* = 8.0 Hz, 2H), 6.95–6.82 (m, 2H), 6.29 (q, *J* = 1.7 Hz, 2H), 4.86 (s, 4H), 4.29 (s, 2H), 3.88 (s, 2H), 3.68–3.36 (m, 8H), 3.13 (t, *J* = 5.2 Hz, 4H), 2.04 (d, *J* = 1.2 Hz, 4H), 1.81 (dd, *J* = 8.9, 1.7 Hz, 1H), 1.64 (d, *J* = 8.8 Hz, 1H), 1.48 (d, *J* = 1.3 Hz, 9H), 1.25 (td, *J* = 7.1, 1.2 Hz, 1H); MS (ESI) calcd for C_42_H_45_N_6_O_5_ (M+H)^+^ 713.3, found 713.4.

To a solution of IWR1-RL (17 mg, 0.024 mmol, 1.0 equiv) in methylene chloride (0.6 mL) was added trifluoroacetic acid (0.3 L). After stirring at 40 °C overnight, the reaction mixture was concentrated and redissolved in acetonitrile (0.5 mL) before [*1323407-86-8*]^77^ (10.8 mg, 0.024 mmol, 1.0 equiv) and *N*,*N*-diisopropylethylamine (13 µL, 0.073 mmol, 3.0 equiv) was added. After stirring at 23 °C overnight, water was added the reaction mixture was extracted with ethyl acetate for three times. The combined organic layers were washed with brine, dried over sodium sulfate, concentrated, and purified by silica gel flash column chromatography to give IWR1-R-OLena (5.5 mg, 23% yield) as a white powder. ^1^H NMR (400 MHz, CD_3_OD) δ 9.00 (d, *J* = 4.5 Hz, 1H), 8.90 (dd, *J* = 7.3, 1.6 Hz, 1H), 8.22–8.03 (m, 2H), 7.83–7.56 (m, 8H), 7.53–7.39 (m, 7H), 7.31 (dd, *J* = 8.1, 0.9 Hz, 1H), 7.16–7.09 (m, 2H), 6.32 (t, *J* = 1.9 Hz, 2H), 5.35 (s, 2H), 5.18 (dd, *J* = 13.3, 5.2 Hz, 1H), 4.64–4.36 (m, 6H), 3.60 (dd, *J* = 3.0, 1.6 Hz, 2H), 3.48 (dq, *J* = 3.4, 1.7 Hz, 2H), 2.94 (ddd, *J* = 18.5, 13.5, 5.5 Hz, 1H), 2.81 (ddd, *J* = 17.6, 4.7, 2.4 Hz, 1H), 2.52 (qd, *J* = 13.2, 4.6 Hz, 1H), 2.21 (dtd, *J* = 12.7, 5.3, 2.5 Hz, 1H), 1.81 (dt, *J* = 8.8, 1.7 Hz, 1H), 1.73 (d, *J* = 8.8 Hz, 1H), 1.32 (s, 1H); MS (ESI) calcd for C_58_H_55_N_8_O_7_ (M+H)^+^ 975.4, found 975.4.

#### Synthesis of G007-LK based PROTACs

Prepared using the same methods as described for the IWR1 based PROTACs.

#### Synthesis of IWR1-PEG3-NCT

To a solution of 6-carboxy-NCT^78^ (4.0 mg, 0.0086 mmol, 1.0 equiv) in *N*,*N*-dimethylformamide (1 mL) were added *N*,*N*-diisopropylethylamine (5 μL, 0.025 mmol, 3.0 equiv) and *N*,*N*,*N*′,*N*′-tetramethyl-*O*-(*N*-succinimidyl)uranium tetrafluoroborate (TSTU, 3.0 mg, 0.0010 mmol, 1.2 equiv). After stirring at 23 °C for 1 h, propargyl PEG3-amine (1.6 mg, 0.0086 mmol, 1.0 equiv) and *N*,*N*-diisopropylethylamine (2 μL, 0.017 mmol, 2.0 equiv) in *N*,*N*-dimethylformamide (0.5 mL) were added. The reaction mixture was stirred at 23 °C for 45 min before concentrated and purified by preparative HPLC to give 6-(propargyl-PEG3-carbamoyl)-NCT (3.5 mg, 56% yield). MS (ESI) calcd for C_44_H_50_N_3_O_7_ (M+H)^+^ 732.4, found 732.3.

To a solution of azido-IWR2 (1.2 mg, 0.0027 mmol, 1.0 equiv) in aqueous acetonitrile (67%, 2.25 mL) was added 6-(propargyl-PEG3-carbamoyl)-NCT (2.0 mg, 0.0027 mmol, 1.1 equiv), copper powder (1.0 mg, 0.0013 mmol, 0.5 equiv). After stirring at 23 °C for 16 h, the reaction mixture was filtered, concentrated, and purified by preparative HPLC to give IWR1-PEG3-NCT (1.0 mg, 31% yield) as a blue solid. ^1^H NMR (600 MHz, MeOD) δ 8.75 (dd, *J* = 15.4, 5.9 Hz, 2H), 8.28 (d, *J* = 8.2 Hz, 1H), 8.11–8.08 (m, 1H), 8.05 (d, *J* = 8.8 Hz, 2H), 7.85 (d, *J* = 8.5 Hz, 1H), 7.78 (s, 1H), 7.57 (t, *J* = 8.1 Hz, 1H), 7.42 (d, *J* = 8.2 Hz, 1H), 7.38 (d, *J* = 8.1 Hz, 1H), 7.07 (d, *J* = 4.4 Hz, 1H), 6.53 (s, 1H), 6.34–6.22 (m, 2H), 6.14 (s, 1H), 5.95 (s, 1H), 5.35 (t, *J* = 5.0 Hz, 1H), 4.58 (s, 2H), 3.86 (q, *J* = 9.2 Hz, 2H), 3.70–3.63 (m, 4H), 3.61 (s, 2H), 3.60–3.56 (m, 4H), 3.49 (t, *J* = 5.8 Hz, 2H), 3.47–3.40 (m, 4H), 3.19 (m, 1H), 2.97–2.85 (m, 4H), 2.75–2.64 (m, 2H), 2.61 (d, *J* = 7.9 Hz, 2H), 2.52–2.43 (m, 4H), 2.28 (t, *J* = 7.4 Hz, 2H), 2.23–2.15 (m, 2H), 2.13 (s, 1H), 2.02 (d, *J* = 13.6 Hz, 4H), 1.91–1.84 (m, 2H), 1.70 (s, 4H), 1.60 (d, *J* = 5.9 Hz, 4H); MS (ESI) calcd for C_70_H_69_N_9_O_10_ (M+H)^+^ 1195.5, found 1196.3.

## Acknowledgements

We thank Dr. Susan Smith (New York University) for providing 293T-TNKS1/2-DKO cells and FLAG-TNKS1-PD construct, Dr. Mariann Bienz and Dr. Melissa Gammons (MRC Laboratory of Molecular Biology) for 293T-DVL2-GFP/AXIN1-dsRed cells and AXIN1-GFP construct, Dr. Dominic Bernkopf (Friedrich-Alexander University Erlangen-Nürnberg) for GFP-AXIN1 and GFP-AXIN2 constructs. We also thank Dr. Katherine Phelps and Dr. Marcel Mettlen for assistance in immunofluorescence microscopy experiments, Dr. Min Fang in automated liquid dispensing and handling, John Shelton in spheroid sectioning, Qing Ding in TMT labeling, and Dr. Duojia Pan, Dr. Xuewu Zhang, Dr. Michael Cohen (Oregon Health & Science University), and Dr. Mike White (IDEAYA Bioscience) for helpful discussions. This work is supported by National Institutes of Health grant R01 CA269377 (to C.C.), R35 GM134883 (to Y.Y.), and P30 CA142543 (to J.W.S.).

## Supplementary Figures

**Figure S1.**
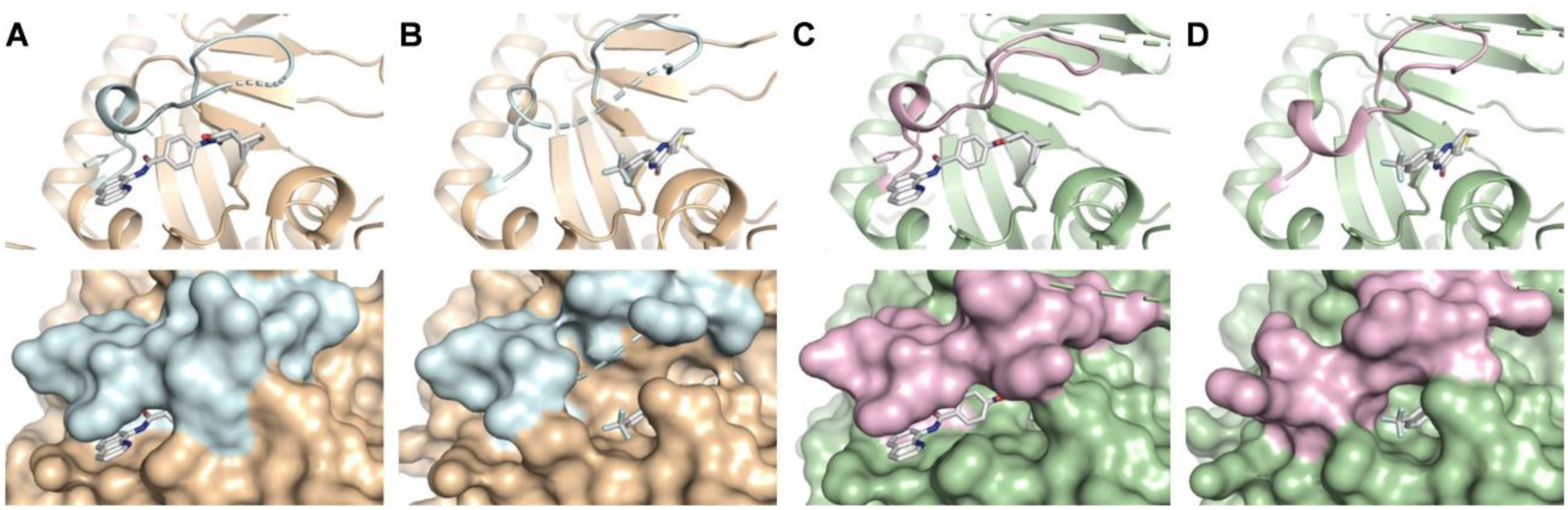
Crystal structures used to guide the design of PROTAC molecules. (A) The crystal structure of TNKS1 with IWR1-exo (PDB 4OA7). (B) The crystal structure of TNKS1 with XAV939 (PDB 3UH4). (C) The crystal structure of TNKS2 with IWR1 (PDB 3UA9). (D) The crystal structure of TNKS2 with XAV939 (PDB 3KR8).

**Figure S2.**
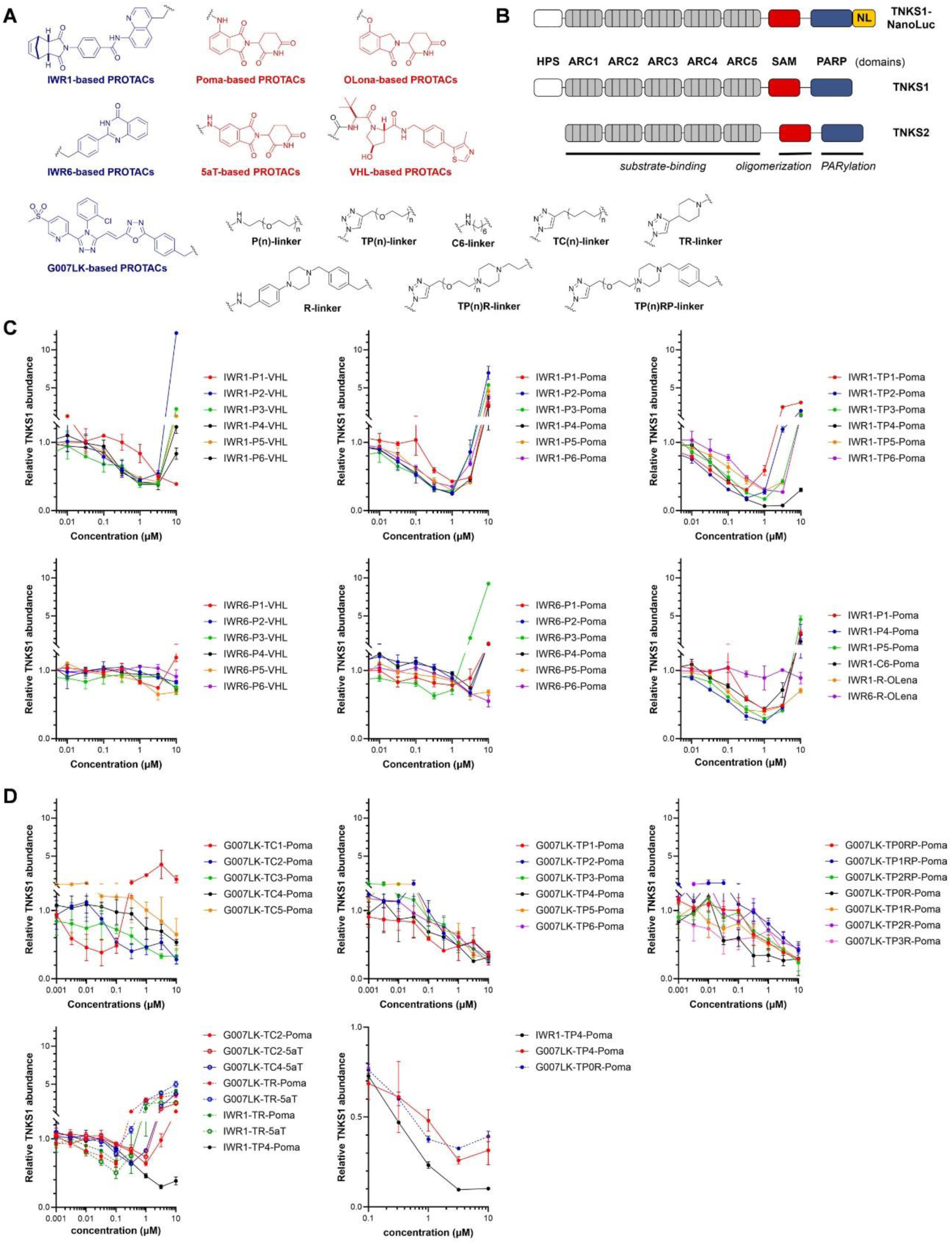
Identification of active PROTAC molecules using CRISPR engineered HAP1 cells expressing a TNKS1-NanoLuc fusion protein. (A) Chemical structures of the PROTAC molecules. (B) Schematic diagrams of the domain structures of TNKS1, TNKS2 and TNKS1-NanoLuc. (C) Relative abundance of the endogenous TNKS1 measured by the luciferase activity upon treating HAP1-TNKS1-NanoLuc cells with IWR1-P(n)-VHL, IWR6-P(n)-VHL, IWR1-P(n)-Poma, IWR6-P(n)-Poma, or IWR1-TP(n)-Poma. IWR1-R-Olena bearing a rigid linker of length and polarity comparable to IWR1-P4-Poma and IWR1-P5-Poma alleviated the hook effect but was less effectively in promoting TNKS1 degradation. Removing the oxygen atom from the linker of IWR1-P1-Poma gave IWR1-C6-Poma with a more hydrophobic linker, but there was no improvement in the degradation efficacy. At high PROTAC concentrations, binding of TNKS and CRBN/VHL by separate PROTAC molecules impedes the formation of productive ternary complexes, resulting in reduced degradation efficacy and consequently the hook effect. Under these conditions, PROTAC molecules function as catalytic inhibitors to prevent TNKS turnover, leading to TNKS accumulation. Data are presented as mean ± SEM (n = 3 biological samples). (D) Selected SAR of PROTACs using G007-LK as the warhead. When shown, IWR1-POMA was included in the assay for direction comparison. All data are presented as mean ± SEM (n = 3 biological samples).

**Figure S3.**
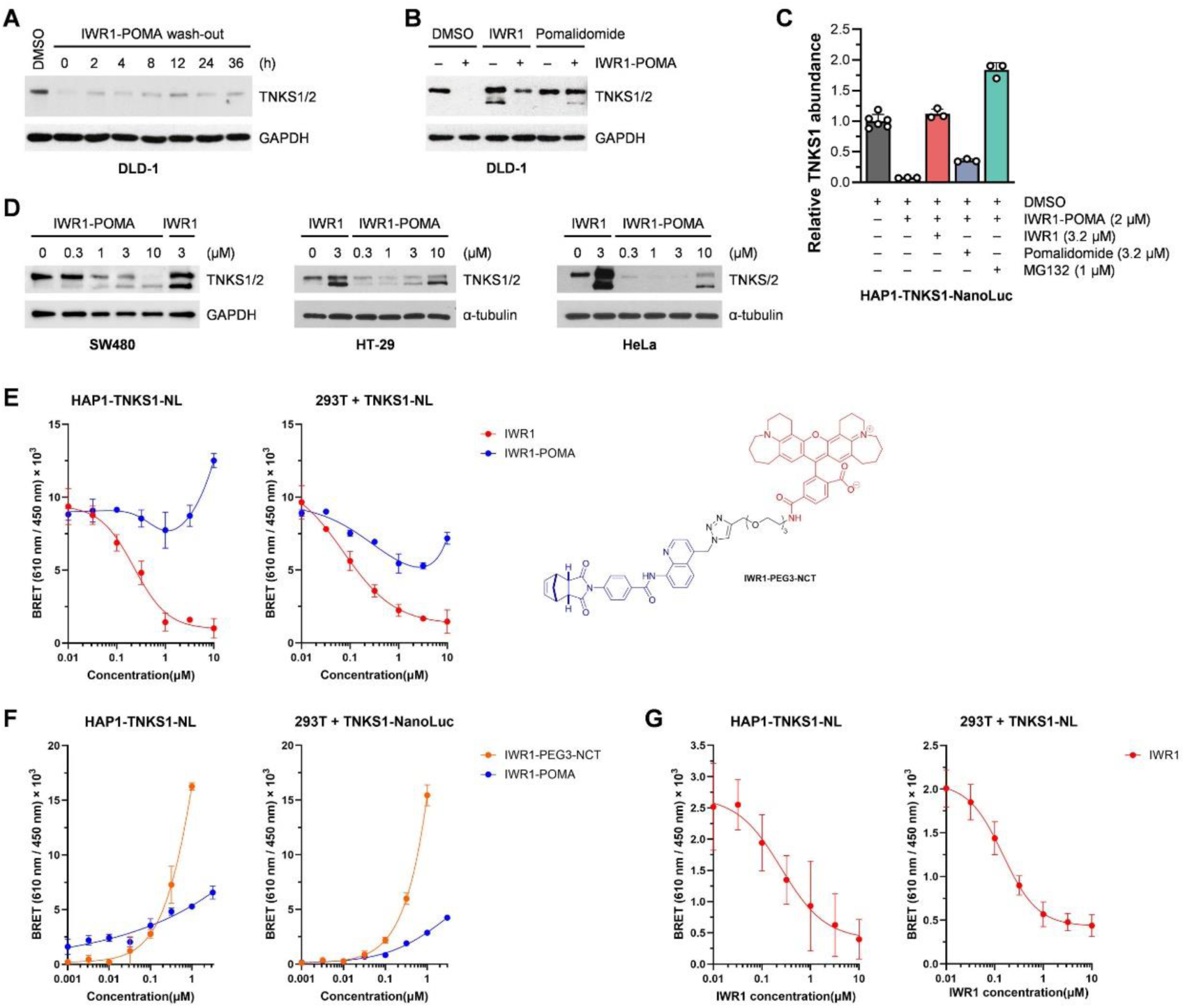
Additional characterization of IWR1-POMA. (A) TNKS did not recover in DLD-1 cells at least 36 h after removing IWR1-POMA. (B) IWR1 (3 µM) and pomalidomide (3 µM) prevented the degradation of TNKS by IWR1-POMA (3 µM) in DLD-1 cells. (C) IWR1, pomalidomide, and MG132 blocked the degradation of TNKS by IWR1-POMA in HAP1-TNKS-NanoLuc cells. (D) IWR1-POMA promoted TNKS degradation while IWR1 induced TNKS accumulation in SW480, HT-29 and HeLa cells. (E) IWR1 suppressed the BRET signals of IWR1-PEG3-NCT (0.5 μM) with an IC50 value of 0.23 μM in HAP1-TNKS1-NanoLuc cells pretreated with MG132 for 16 h or with an IC50 value of 0.07 μM in 293T cells transfected with TNKS1-NanoLuc plasmid (25 ng). However, the intrinsic fluorescence of IWR1-POMA prevented the determination of its binding to TNKS1 by this method. Data are presented as mean ± SEM (n = 3 biological samples). (F) Addition of IWR1-PEG3-NCT or IWR1-POMA increased BRET signals in a dose-dependent manner in HAP1-TNKS1-NanoLuc cells pretreated with MG132 for 16 h or in 293T cells transfected with TNKS1-NanoLuc plasmid (25 ng). Data are presented as mean ± SEM (n = 3 biological samples). (G) IWR1 suppressed the BRET signals of IWR1-POMA (0.5 μM) with an IC50 value of 0.25 μM in HAP1-TNKS1-NanoLuc cells pretreated with MG132 for 16 h or with an IC50 value of 0.15 μM in 293T cells transfected with TNKS1-NanoLuc plasmid (25 ng). Data are presented as mean ± SEM (n = 6 biological samples).

**Figure S4.**
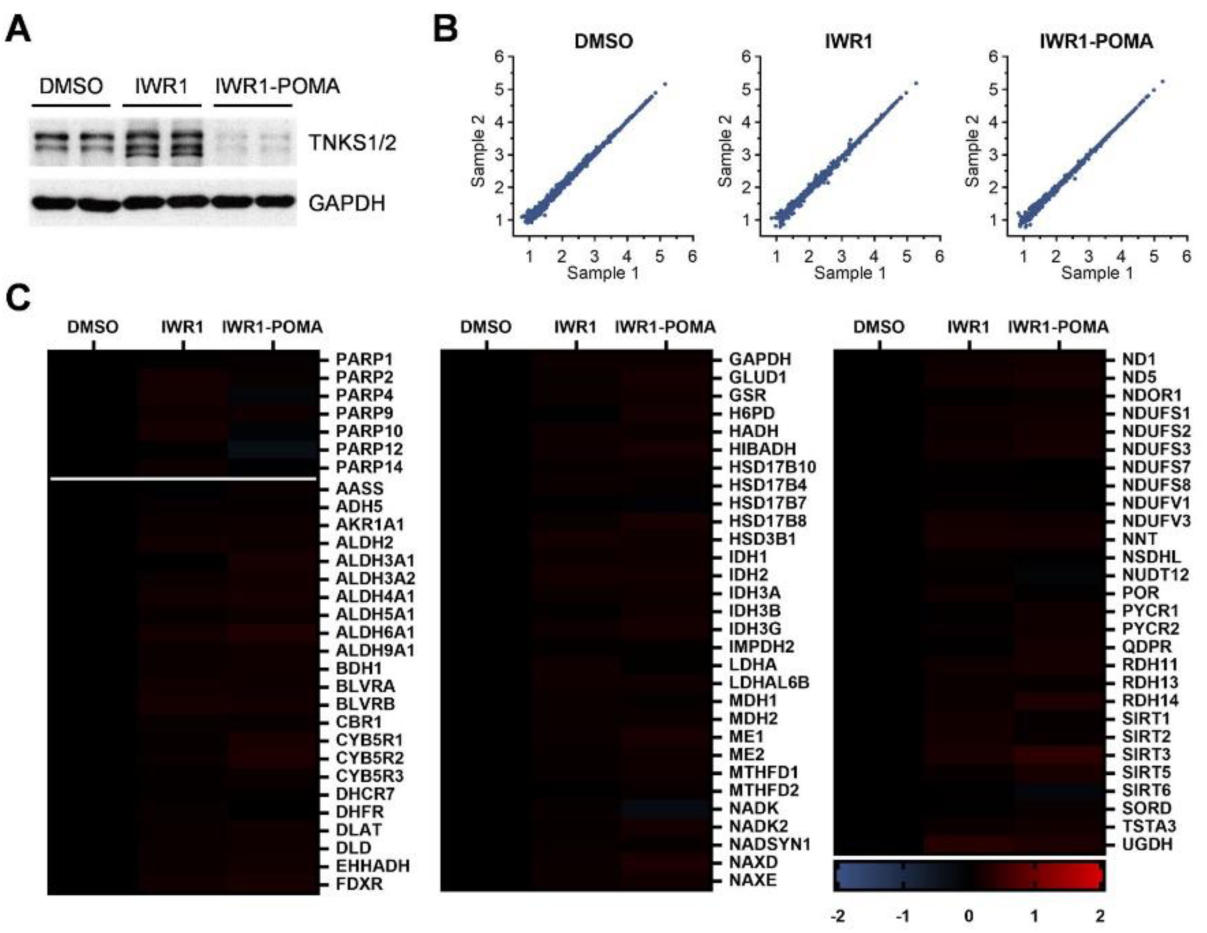
Proteomic analysis of DLD-1 cells treated with DMSO, IWR1 or IWR1-POMA. (A) Western blot analysis of samples corresponding to Figure 1 E and 1F confirmed the accumulation of TNKS1/2 by IWR1 and the depletion by IWR1-POMA. (B) Correlation analysis showed high reproducibility between the two biological samples. (C) IWR1-POMA selectively degraded TNKS without inducing appreciable perturbations to 7 other PARP family member proteins and 79 NAD(P)-dependent enzymes detected in this proteomic experiment.

**Figure S5.**
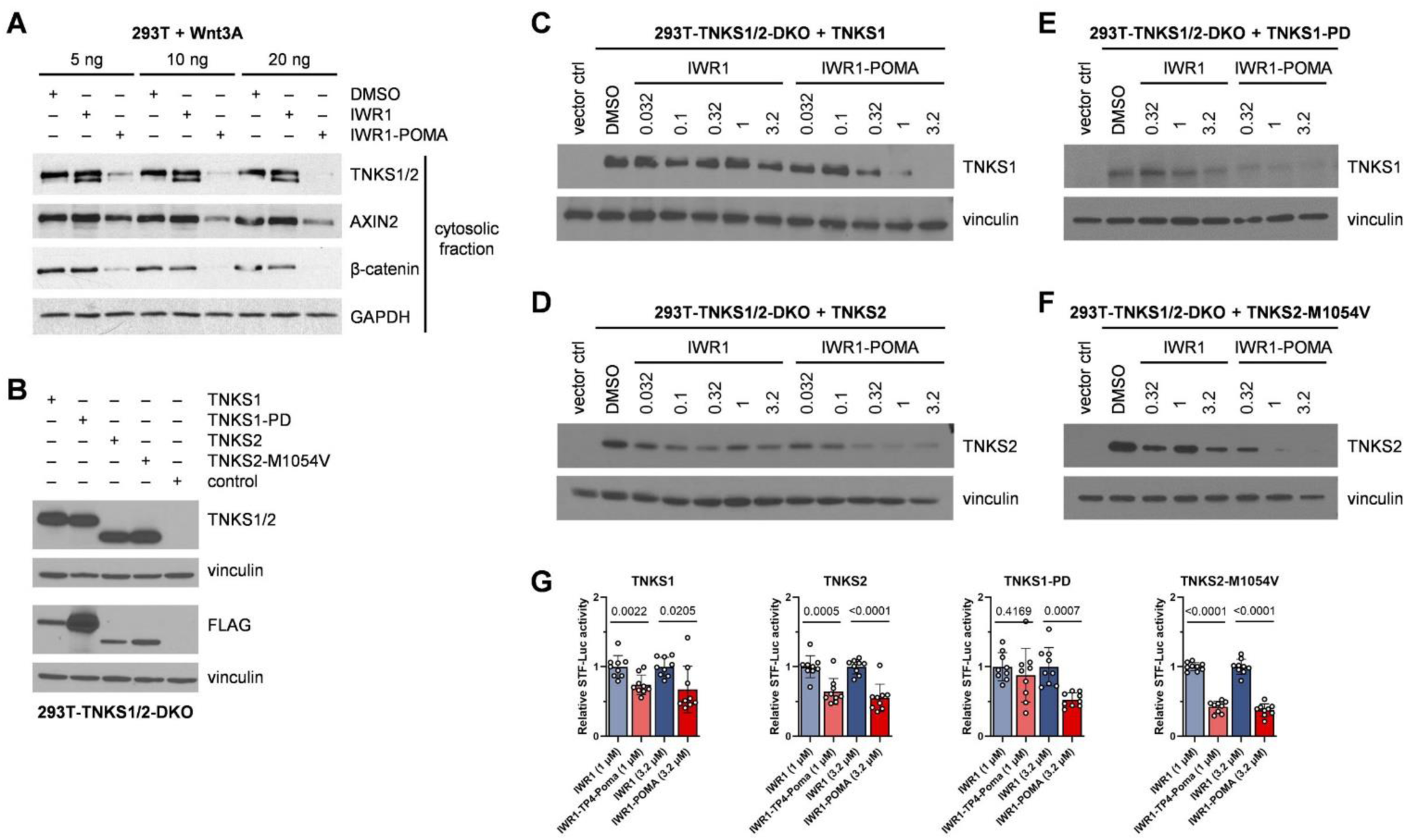
Degradation of TNKS by IWR1-POMA in 293T cells. (A) 293T cells were transfected with different doses of Wnt3A plasmid and then treated with DMSO, IWR1 (3 μM) or IWR1-POMA (3 μM). The cytosolic fraction of the cell lysates was then examined by Western blot. IWR1-POMA promoted a more complete degradation of β-catenin than IWR1 and reduced the level of the WNT target AXIN2. (B) Western blot analysis confirmed the lack of TNKS expression in 293T-TNKS1/2-DKO cells and validated the expression of FLAG-TNKS1, 3×FLAG-TNKS1-PD, FLAG-TNKS2 and FLAG-TNKS2-M1054V after transfection. (C) Western blot analysis of samples corresponding to Figure 2A confirmed the degradation of TNKS1 by IWR1-POMA. (D) Western blot analysis of samples corresponding to Figure 2B confirmed the degradation of TNKS2 by IWR1-POMA. (E) Western blot analysis of samples corresponding to Figure 2C confirmed the degradation of TNKS1-PD by IWR1-POMA. (F) Western blot analysis of samples corresponding to Figure 2D confirmed the degradation of TNKS2-M1054V by IWR1-POMA. (G) Comparison of the effects of IWR1 and IWR1-POMA on WNT/β-catenin pathway activity in 293T-TNKS1/2-DKO cells, related to Figure 2. Data were normalized to the STF activity of the IWR1-treated samples at the indicated concentrations to combine results from three independent experiments for statistical analysis. Data are presented as mean ± SEM (n = 3×3 biological samples) with p-values calculated by two-tailed unpaired t-test.

**Figure S6.**
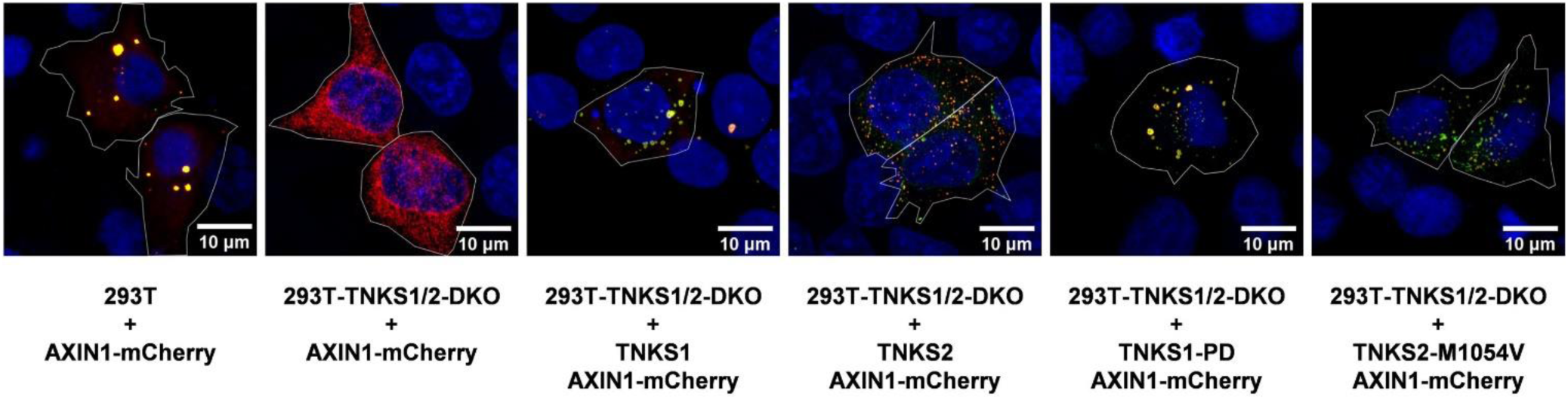
Effects of TNKS on AXIN puncta. Additional images of 293T or 293T-TNKS1/2-DKO cells transfected with or without TNKS plasmids from independent experiments related to Figure 3.

**Figure S7.**
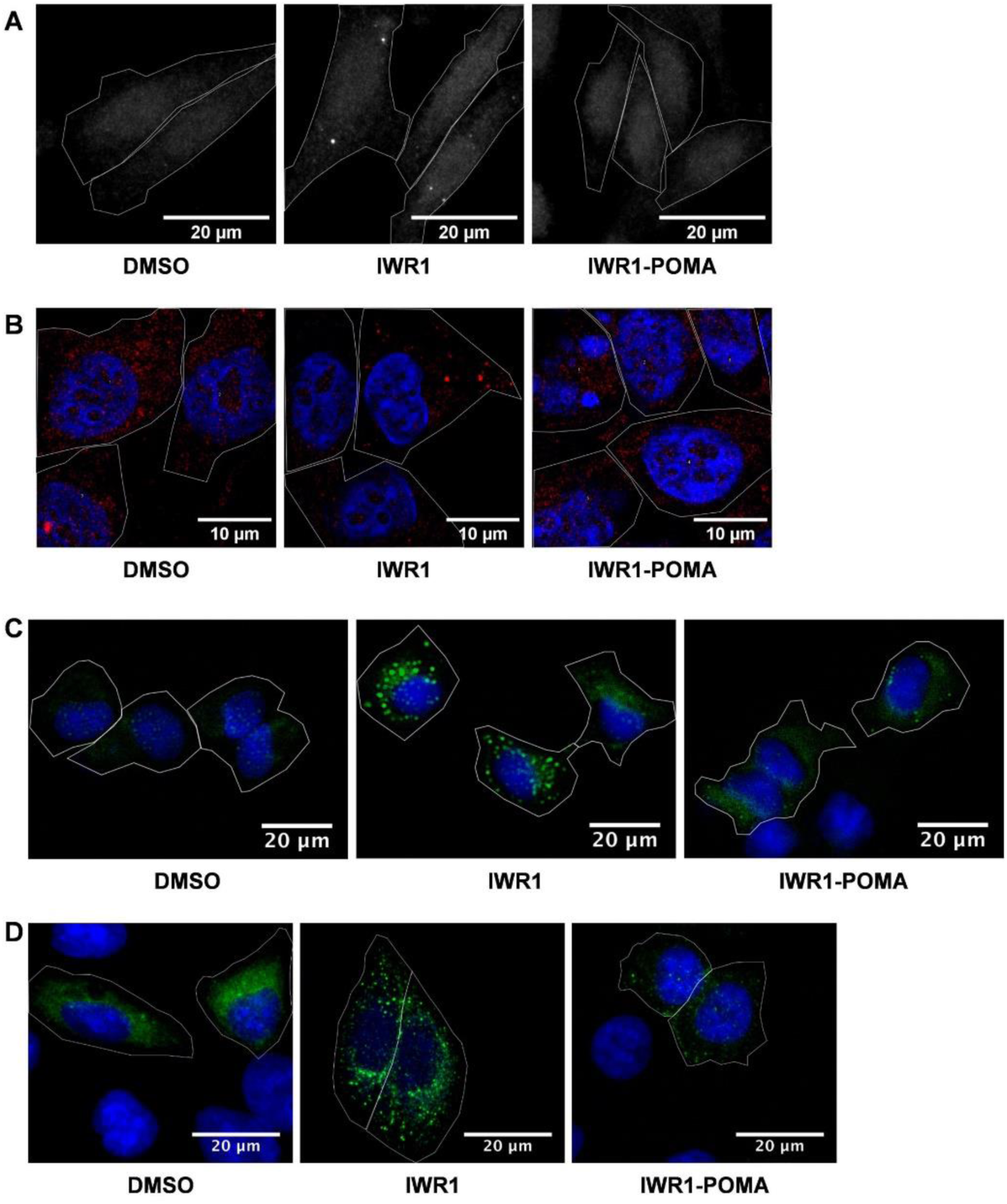
TNKSi induced the formation of AXIN puncta. (A) Additional images of SW480 cells treated with DMSO, IWR1 (5 μM), or IWR1-POMA (1 μM), related to Figure 4A. (B) Multi-channel images of HEK293-AXIN1-dsRed-KI cells treated with DMSO, IWR1 (3 μM), or IWR1-POMA (3 μM) with nuclear staining (DAPI, blue), corresponding to Figure 4C. (C) HeLa cells transfected with GFP-AXIN1 plasmid followed by treating with DMSO, IWR1 (3 µM), or IWR1-POMA (3 µM). Together with Figure 4C, this experiment shows that the position of GFP tag does not affect puncta formation. (D) HeLa cells transfected with GFP-AXIN2 plasmid followed by treating with DMSO, IWR1 (3 µM), or IWR1-POMA (3 µM). This experiment shows that AXIN2 also forms puncta upon IWR1 treatment.

**Figure S8.**
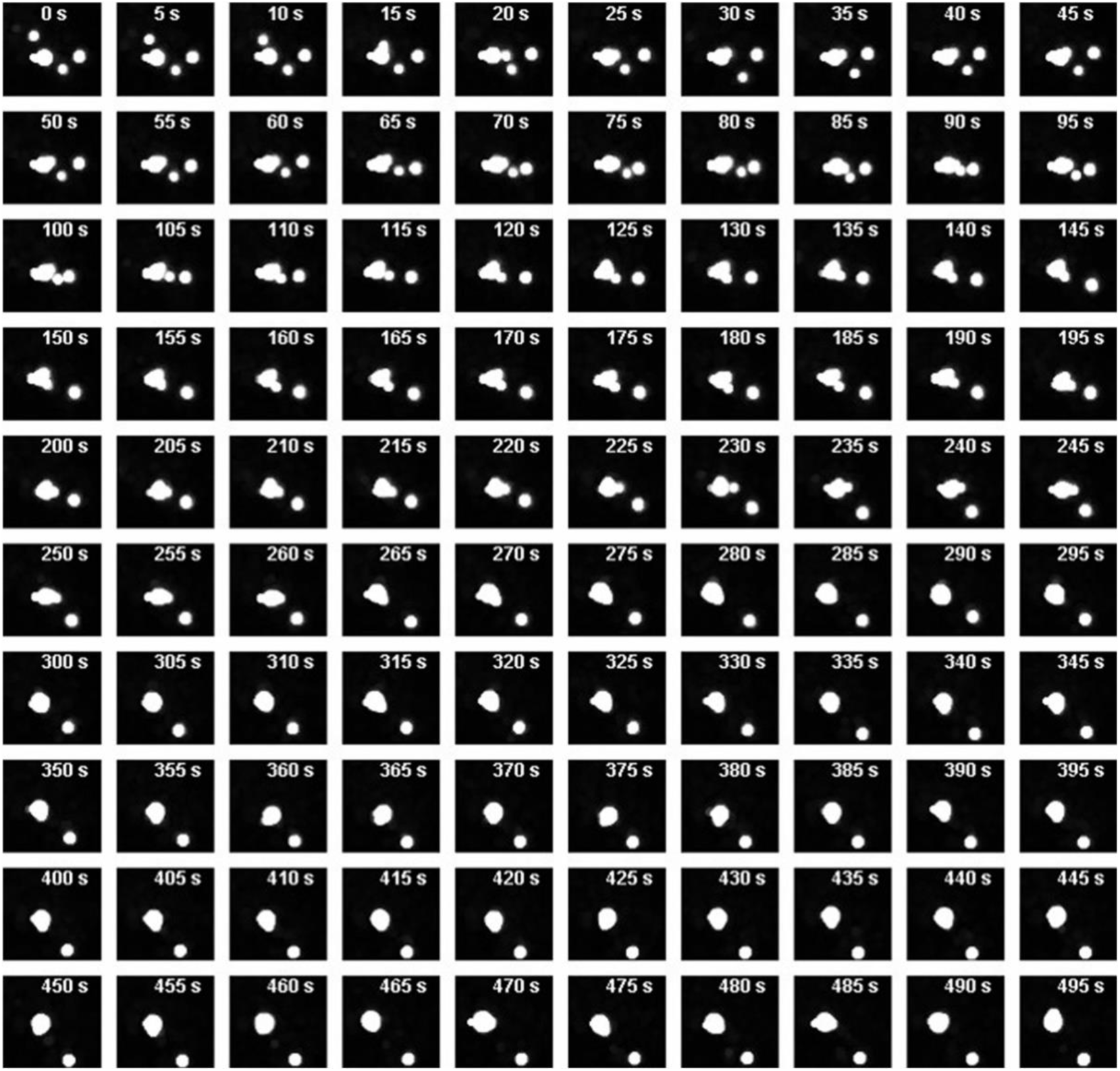
Time-lapse imaging supports the liquid-like properties of AXIN puncta. U2OS cells transfected with AXIN1-GFP plasmid led to the formation of AXIN puncta that underwent dynamic fusion upon contact with transient budding-like shape fluctuations.

**Figure S9.**
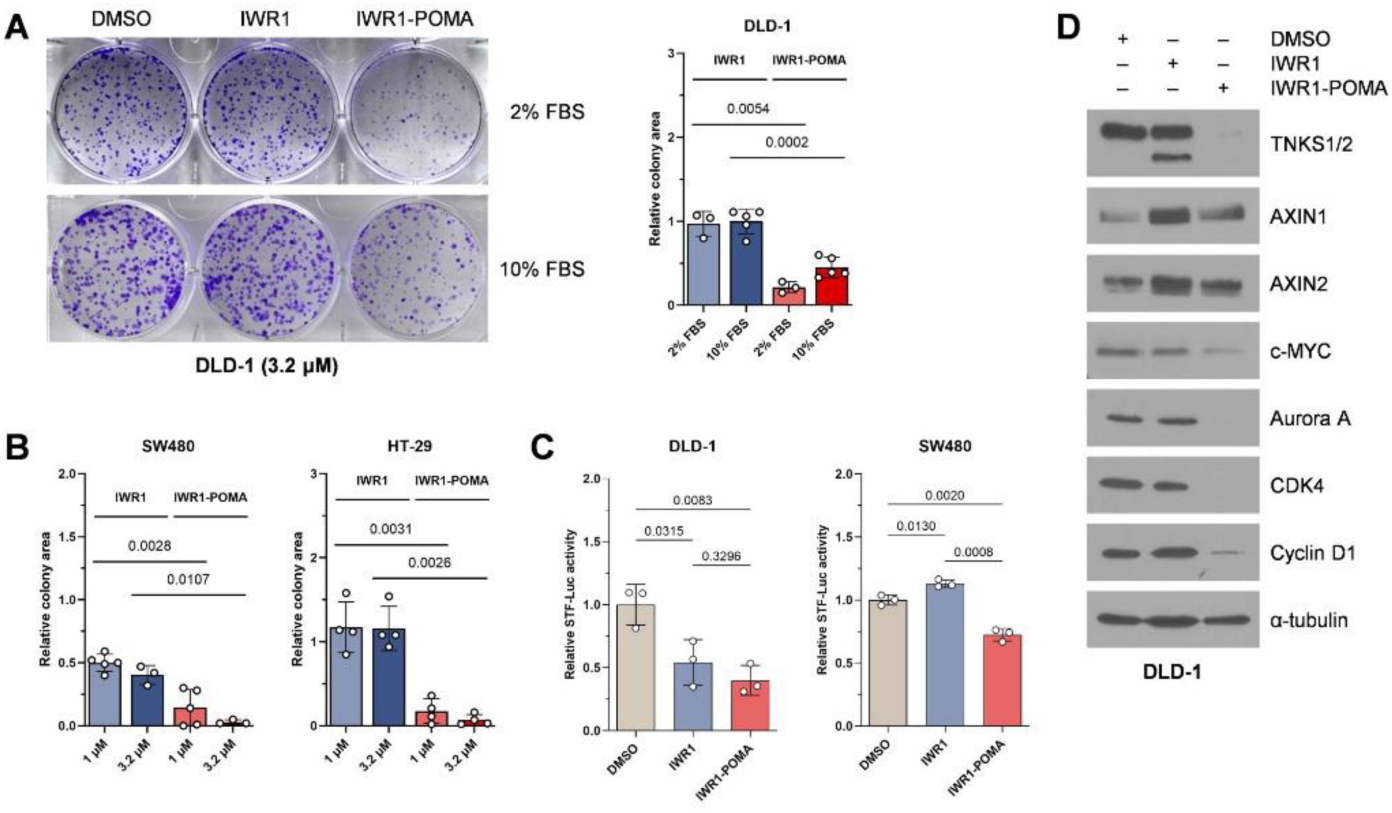
IWR1-POMA suppressed CRC growth through WNT inhibition. (A) IWR1-POMA suppressed the formation of DLD-1 colonies more effectively than IWR1 under both normal and low serum conditions. Data are presented as mean ± SEM (n = 3–4 biological samples) with p-values calculated by two-tailed unpaired t-test. (B) Quantification of SW480 and HT-29 colonies, related to Figure 6A. Data are presented as mean ± SEM (n = 3–5 biological samples) with p-values calculated by two-tailed unpaired t-test. (C) IWR1-POMA (3 μM) suppressed of WNT signaling more effectively than IWR1 (3 μM) in DLD-1 and SW480 cells. Data are presented as mean ± SEM (n = 3 biological samples) with p-values calculated by two-tailed unpaired t-test. (D) Western blot analysis confirmed that c-MYC, Aurora A, CDK4 and cyclin D1 responded to IWR1-POMA (3 μM) but not IWR1 (3 μM) in DLD-1 cells.

**Figure S10.**
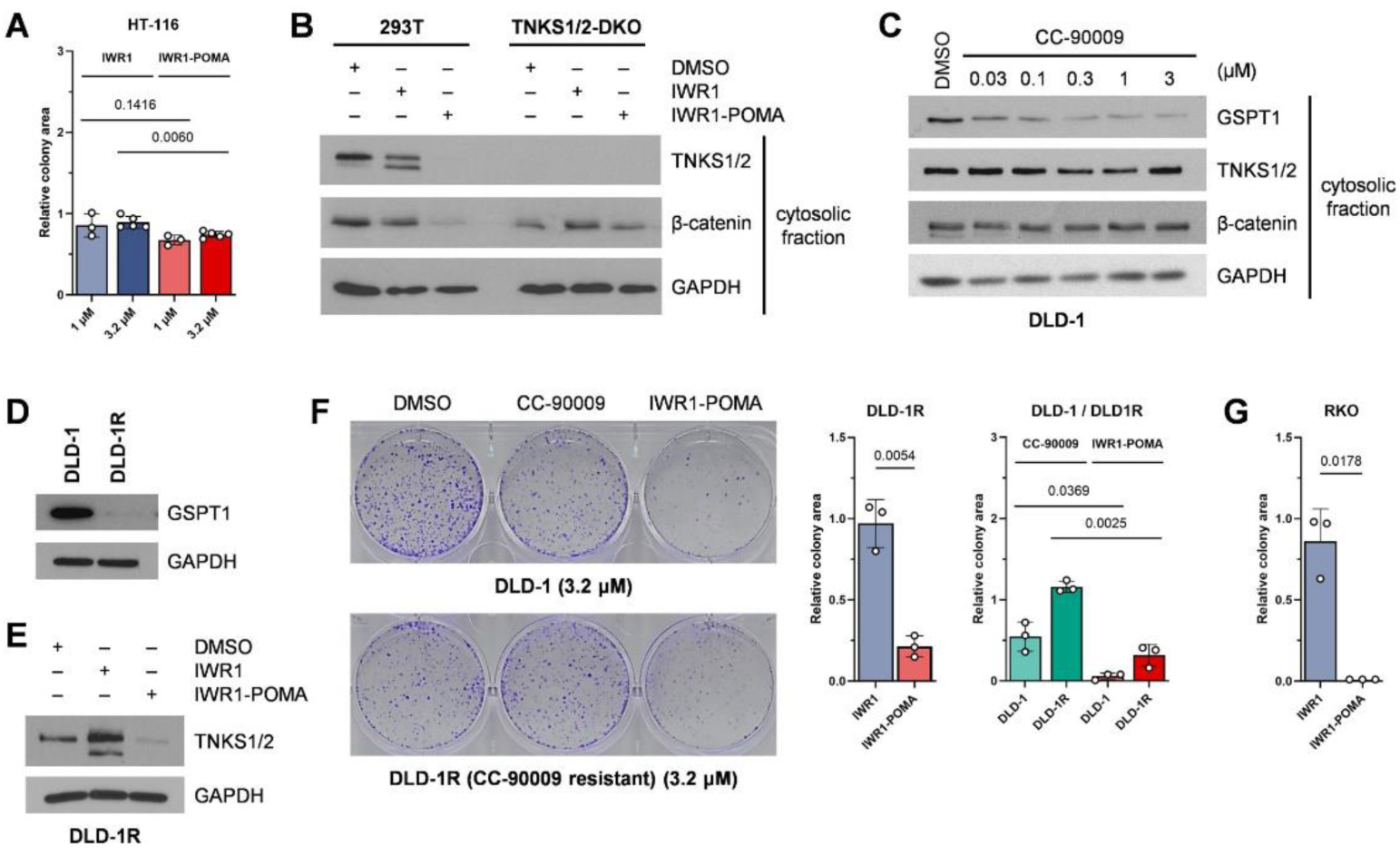
IWR1-POMA suppressed CRC proliferation through on-target WNT inhibition. (A) Quantification of HCT116 colonies, related to Figure 6B. Data are presented as mean ± SEM (n = 3–4 biological samples). (B) IWR1-POMA (3 µM) reduced the cytosolic β-catenin level more effectively than IWR1 (3 µM) in 293T cells. The level of cytosolic β-catenin in 293T-TNKS1/2-DKO cells that lack both TNKS1 and TNKS2 did not change significantly with either drug treatment. (C) CC-90009 induced GSPT1 degradation in DLD-1 cells but had little effect on TNKS. GSPT2 was not detectable by Western blot, which is consistent with the reported GSPT levels determined by quantitative proteomic analysis in this cell line—102,567 ppb for GSPT1 and 2,495 ppb for GSPT2 (https://www.ebi.ac.uk/gxa/experiments/E-PROT-18/Results)^69^. (D) DLD-1R cells obtained from cultivating DLD-1 cells with CC-90009 have a dramatically reduced level of GSPT1 expression meanwhile maintaining normal TNKS expression. (E) IWR1 (3 μM) promoted TNKS accumulation while IWR1-POMA (3 μM) induced TNKS degradation in DLD-1R cells. (F) IWR1-POMA prevented colony formation of DLD-1R cells deficient in GSPT1/2. Data are presented as mean ± SEM (n = 3 biological samples) with p-values calculated by two-tailed unpaired t-test. (G) Quantification of RKO colonies, related to Figure 6B. Data are presented as mean ± SEM (n = 3 biological samples) with p-values calculated by two-tailed unpaired t-test.

**Figure S11.**
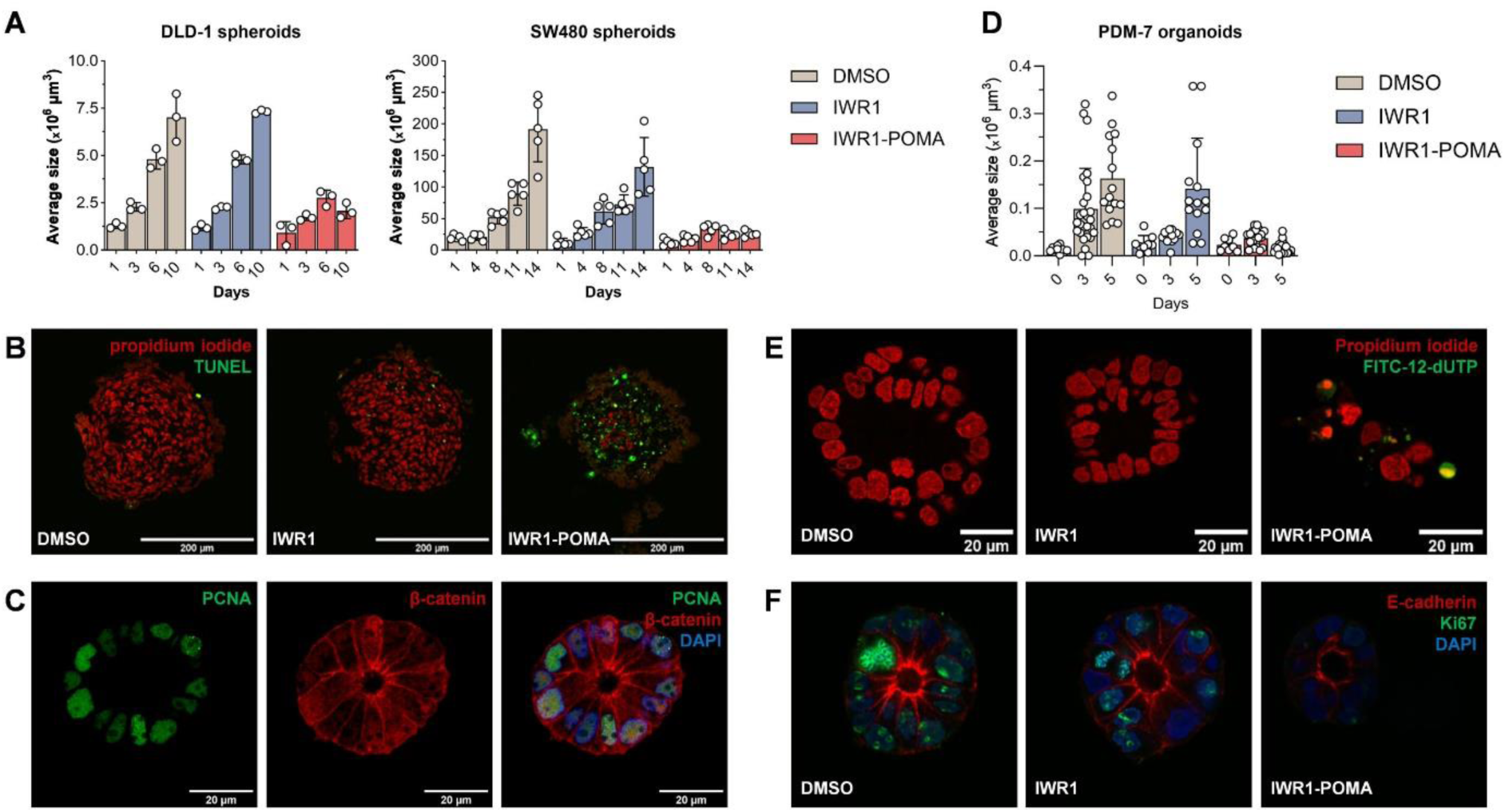
IWR1-POMA demonstrated efficacy in CRC spheroid and primary organoid models. (A) The growth chart of DLD-1 and SW480 spheroids treated with DMSO, IWR1 (5 μM), or IWR1-POMA (5 μM). The sizes represent the apparent dimensions of the spheroids including the peripheral dead cells. Data are presented as mean ± SEM (n = 3 or 5 biological samples) with p-values calculated by two-tailed unpaired t-test. (B) IWR1-POMA (5 µM) induced apoptosis in DLD-1 spheroids. (C) PDM-7 organoids grown from single cells preserved the heterogeneous nature of CRC tumors. (D) IWR1-POMA (1 μM) suppressed the formation of PDM-7 organoids from single cells while IWR1 (1 μM) did not. Data are presented as mean ± SEM (n = 8–27 biological samples) with p-values calculated by two-tailed unpaired t-test. (E and F) IWR1-POMA (1 μM) suppressed proliferation and induced apoptosis in PDM-7 organoids grown from single cells while IWR1 (1 μM) had little effect.

